# scELMo: Embeddings from Language Models are Good Learners for Single-cell Data Analysis

**DOI:** 10.1101/2023.12.07.569910

**Authors:** Tianyu Liu, Tianqi Chen, Wangjie Zheng, Xiao Luo, Yiqun Chen, Hongyu Zhao

**Affiliations:** Interdepartmental Program in Computational Biology & Bioinformatics, Yale University, New Haven, 06511, CT, USA; Department of Biostatistics, Yale University, New Haven, 06511, CT, USA; Department of Computer Science, University of California, Los Angeles, Los Angeles, 90095, CA, USA; Department of Biostatistics, Johns Hopkins University, Baltimore, 21218, MD, USA

**Keywords:** Single-Cell Data Analysis, Foundation Model, Large Language Model, In-silico Treatment Analysis, Perturbation Analysis

## Abstract

Various Foundation Models (FMs) have been built based on the pre-training and fine-tuning framework to analyze single-cell data with different degrees of success. In this manuscript, we propose a method named scELMo (Single-cell Embedding from Language Models), to analyze single-cell data that utilizes Large Language Models (LLMs) as a generator for both the description of metadata information and the embeddings for such descriptions. We combine the embeddings from LLMs with the raw data under the zero-shot learning framework to further extend its function by using the fine-tuning framework to handle different tasks. We demonstrate that scELMo is capable of cell clustering, batch effect correction, and cell-type annotation without training a new model. Moreover, the fine-tuning framework of scELMo can help with more challenging tasks including in-silico treatment analysis or modeling perturbation. scELMo has a lighter structure and lower requirements for resources. Our method also outperforms recent large-scale FMs (such as scGPT [1], Geneformer [2]) and other LLM-based single-cell data analysis pipelines (such as GenePT [3] and GPTCelltype [4]) based on our evaluations, suggesting a promising path for developing domain-specific FMs.

## 1 Introduction

Developing foundation models (FMs) has become critical for broad areas including but not restricted to engineering and sciences [5–7], and large language models (LLMs) are examples of successful FMs. In biology, FMs have been developed to analyze DNA sequences [8, 9], represent cells and genes [1, 2, 10, 11], and perform other tasks. It has been shown that FMs can facilitate various downstream biological analyses. Here we focus on a specific type of biomedical tabular data, known as single-cell sequencing data [12, 13], for its intersection with FMs. Single-cell sequencing data can describe biological activity and information at the cell level. The units are individual cells, and the features can be gene expression levels [13], protein expression levels [14, 15], methylation levels [16], and others.

Several pre-training-based FMs have been developed to analyze single-cell data based on large-scale sequencing datasets collected from different studies. For example, scGPT [1], Geneformer [2], and scFoundation [17] are pre-trained with large-scale single-cell transcriptomics to learn the patterns in the data and handle different tasks by either using the cell embeddings/gene embeddings as representations or fine-tuning the pre-trained models with new data. However, these methods require a large amount of computational resources and storage space to complete pre-training, and can be affected by both data pre-processing and gene selection methods. Several analyses [18, 19] have discussed the limitations of these single-cell foundation models by performing comprehensive benchmarking analyses. Meanwhile, the developers of GenePT [3] discussed the limitation of only using the gene expression information for biological analysis, and conjecture that we can either use the information from the National Center for Biotechnology Information (NCBI) [20] as prompts to get the embeddings of such prompts from LLMs to describe the genes (GenePT-w), or transfer cells to sentences by gene ranking and use the ranking results as a prompt to extract the cell embeddings from LLMs (GenePT-s). The results from GenePT can be further explored by incorporating prior knowledge of the features in single-cell datasets. However, GenePT has limited generation ability by relying on external knowledge from NCBI. Moreover, the NCBI database may not be informative enough to summarize the gene functions in a well-structured manner. On the other hand, using GenePTs to generate cell embeddings is not very practical because of the high sparsity of single-cell data, so the genes with zero expression are hard to rank. GenePT-s is also not capable of handling large-scale datasets because of the usage and time limitation from OpenAI API [21]. GenePT also discussed the zero-shot learning ability of such embeddings, but it could not incorporate cell-level metadata information, such as cell types, which further restricted its applications for more tasks. Recently, the access of LLMs including GPT 3.5 [22], GPT 4 [23], and LLaMa [24] offers us opportunities to explore the information of these features with the help of LLMs. These LLMs have been widely used to summarize knowledge [25], design network searching/models [26], and perform other tasks to enhance scientific research [27, 28].

In this manuscript, we explore the ability of using LLMs in a different manner. We generate meaningful text descriptions of cell-level or feature-level metadata as well as embeddings of such descriptions based on LLMs. We assume that the embeddings from LLMs carry biological properties and can be utilized in various downstream applications. Here we introduce scELMo as a pipeline for analyzing single-cell multiomic data based on the text description and embeddings directly from LLMs [29]. Using gene as one example, we leverage LLMs to summarize the functional information of a given gene with a suitable prompt and also use the same LLM to extract the embeddings of such description. We then either incorporate the embeddings directly into the single-cell data by matrix operation or combine the embeddings with other models with fine-tuning targets for various tasks. Different from traditional single-cell Foundation Models, scELMo does not require pre-training with LLM embeddings and saves resources to accelerate biological discoveries and insight validation. We demonstrate that scELMo is a simple but effective tool for single-cell data analysis under both the zero-shot learning framework and the fine-tuning framework, supported by comprehensive benchmarking analyses.

## 2 Results

### Overview of scELMo

The embeddings from LLMs are powerful in preserving the context and information similarity of the original inputs [30–32], and thus they are good priors as a start for incorporating the biological knowledge from enriched text to analyze a specific dataset. Therefore, the high-level idea of scELMo is to transfer the information of each cell from the sequencing data space to the embedding space from LLM. We complete this transformation by incorporating information from the feature space (for example, genes and proteins.) or the cell space (for example, cell types or cell states). To transfer the feature information into the LLM embedded space, we can either use prior knowledge from a known database such as NCBI or the summary given by LLMs as a prompt, and then generate the embeddings for the prompt based on LLMs’ embedding layers. Here we choose GPT 3.5 to summarize the functions of features and to generate embeddings based on our evaluation in the Results section.

After collecting embeddings from LLMs and transcriptomic profiles from cells, we can use them based on the zero-shot learning framework to produce cell embeddings after integration. These cell embeddings can be used for clustering or correcting the batch effect. Such framework means that we utilize matrix operation to integrate the LLM-produced embeddings with transcriptomic profiles to generate new cell embeddings. As several methods demonstrate the improvement of considering biological knowledge in modeling single-cell data [1, 33, 34], we can also combine the embeddings with task-specific models to improve their performance for various downstream tasks, known as the fine-tuning framework which requires training task-specific models again with LLM-produced embeddings as new inputs. Such framework provides a more flexible approach for utilizing gene embeddings as information-enriched biological priors to enhance task-specific methods in downstream applications. Figure 1 shows the flowchart of scELMo under two different frameworks. The details of scELMo under different frameworks are given in the Methods section.

**Fig. 1.**
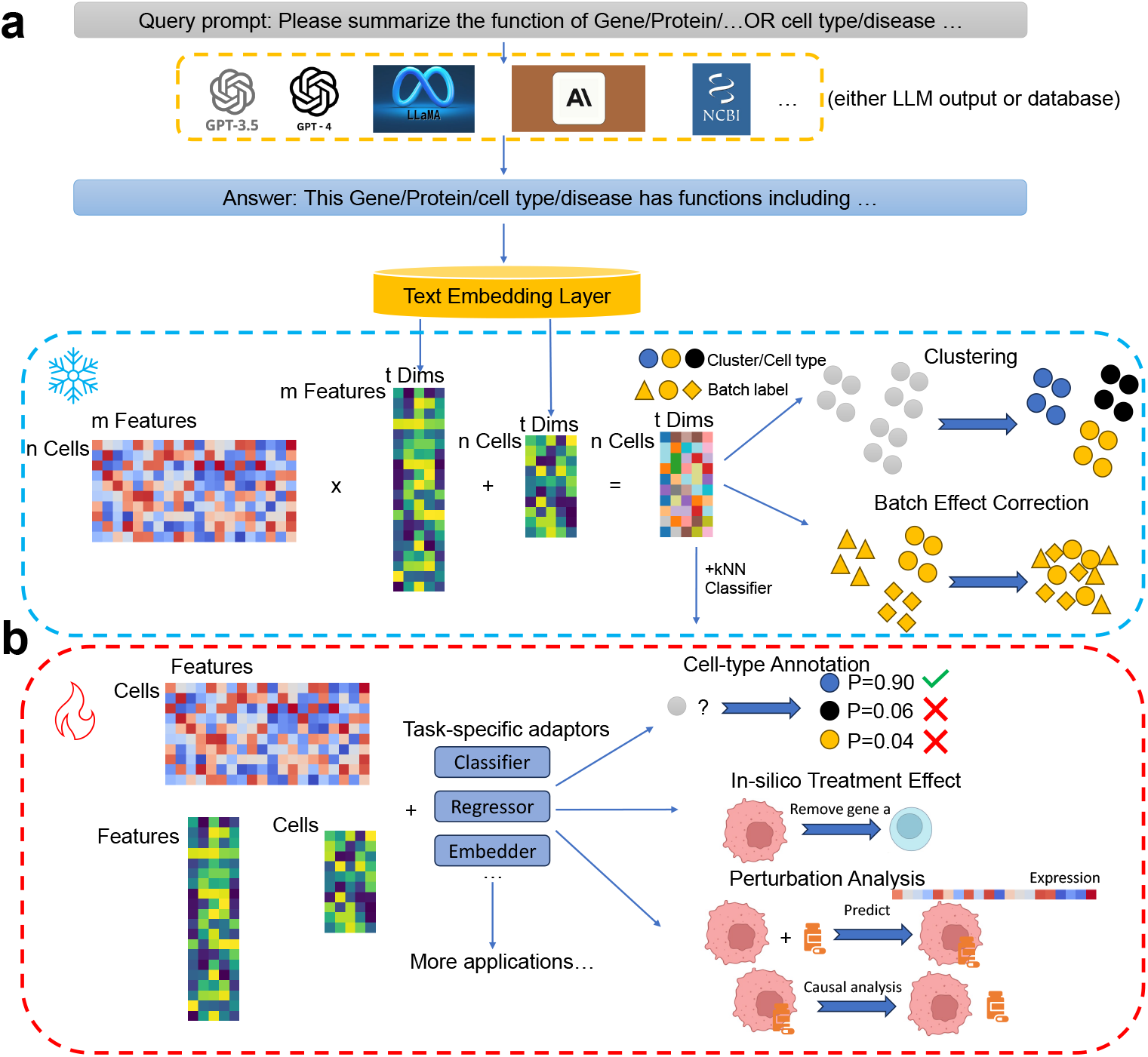
Workflow of scELMo. (a) Zero-shot learning (denoted as ice) framework of scELMo. We extract the text description of metadata by using either databases or LLMs. The prompts are adjustable. We use GPT 3.5 to generate the embeddings of the text descriptions as the embeddings of features (including genes, peaks, etc. which are biological features from cell profiles) or cell states. We then aggregate these embeddings with single-cell profiles to generate cell embeddings. (b) Fine-tuning (denoted as fire) framework of scELMo. We combine embeddings of metadata and single-cell profiles with task-specific adaptors and train the adaptors to address downstream applications.

### Evaluation of Hallucinations in LLMs

The first step of our work is to define a suitable LLM for generating text descriptions and embeddings. A suitable LLM should not suffer from Hallucinations [35], that is, generating fake or incorrect information about our given feature information or cell information. Here we consider GPT 2 [36], GPT 3.5, GPT 4, Llama-2 (70B), Mistral [37], bioGPT [38], Claude 2 [39], and Bard (PaLM 2) [40] as candidates to generate text description. Considering the diversity of tokens as well as the access to embedding layers, we chose GPT 3.5 to generate embeddings. We randomly sampled 20 proteins and 20 genes from known proteins (*∼* 200 proteins from [41]) and genes (*∼* 30,000 genes from NCBI), and used these LLMs we mentioned above to generate the text description of these features. We evaluated the correctness of the outputs by comparing them with known databases (GeneCards [42] and NCBI) based on both Bilingual Evaluation Understudy (BLEU) [43] score and Human-Eval [44] score. BLEU score essentially measures the similarity between the LLM output and the human-created references, focusing on the overlap of n-grams (sequences of n words). The results of genes and proteins are shown in Figure 2 (a), together with the time usage for one query for each LLM, shown in Figure 2 (b). We also include the results of cell types in Extended Data Figure 1 (a).

**Fig. 2.**
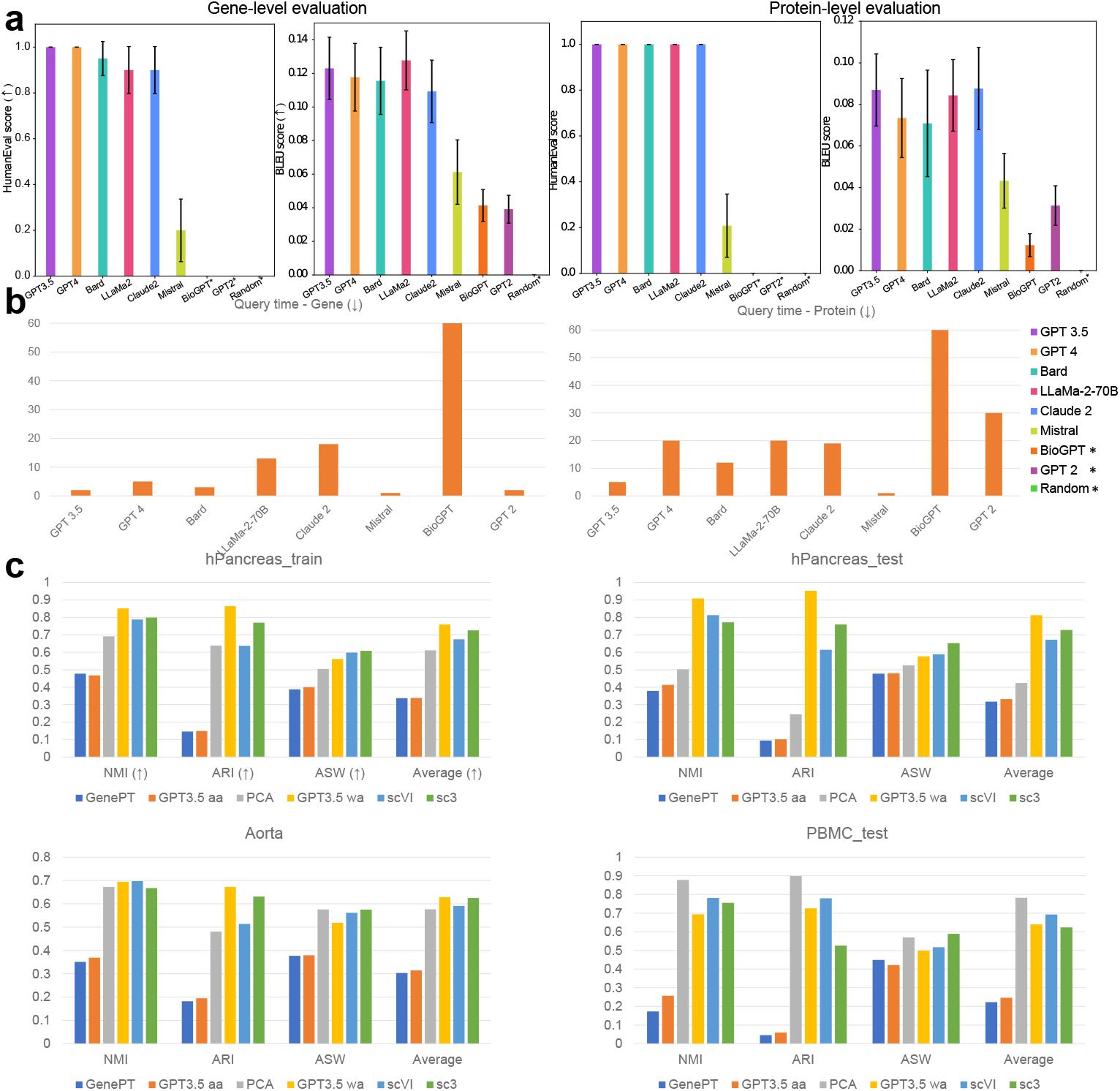
Evaluations of the outputs of LLMs and the clustering performance. *: This sign represents that the selected method has zero Human-Eval score. (a) Metrics for evaluating meaningful outputs of biological features across different LLMs. The left panel represents the BLEU score and Human-Eval score of genes, while the right panel represents the BLEU score and Human-Eval score of proteins. (b) Average query time for each LLM. The left panel represents the query time of genes, and the right panel represents the query time of proteins. (c) Evaluations of the clustering performance based on major methods. Different panels represent the results of different datasets.

Based on our results, GPT 3.5 and GPT 4 gave us the most accurate text description. However, comparing with GPT 3.5, GPT 4 required longer query time and some of the outputs from GPT 4 did not follow our required format mentioned in the prompt. Meanwhile, we also considered analyzing the effects of different prompts as prompt engineering for querying gene information. We also analyzed the results based on MetaPrompt [45] and Chain-of-Thought (COT) [46]. Using MetaPrompt or COT prompting does not lead to improved BLEU scores, with values remaining comparable to the original prompt, shown in Extended Data Figure 1 (b). Moreover, the Human-Eval scores decrease when using these methods compared to the original prompts, shown in the same figure. To evaluate different LLMs more comprehensively, we scaled the evaluation to 100 random genes to compute the BLEU scores between LLM-generated descriptions and NCBI information. Our results show that GPT 3.5 also has the highest BLEU score with low standard error under the scaling evaluation case, followed by Claude, shown in Extended Data Figure 2. By considering the trade-off between query time and quality of generated descriptions, we selected GPT 3.5 as our tool to generate text descriptions of features based on our default prompts. Extended Data Figure 1 (a) also shows that GPT 3.5 is also good at generating descriptions for cell types. Furthermore, we also evaluated the stability of the embeddings of LLM’s outputs based on the same prompt for the same gene, as well as the similarity of embeddings from different genes. Our results are summarized in Extended Data Figures 3 (a) and (b). According to Extended Data Figure 3 (a), the cosine similarity between the embeddings from different LLM outputs of the same gene was higher than 0.9, suggesting that the embeddings from GPT 3.5 have high stability and may capture the intrinsic functions for each gene. Based on Extended Data Figure 3 (b), the embeddings from different genes had lower cosine similarity ([0.74, 0.84]) comparing with the embeddings from the same gene ([0.88, 0.99]). Therefore, the embeddings from GPT 3.5 may also capture the functional heterogeneity of different genes. Moreover, we analyze the effects of gene embeddings from different random seeds on downstream applications in the Discussion section. Furthermore, we loaded the gene functional annotation provided by Geneformer [2] to predict the functional information for the other genes based on a k-nearest-neighbor (kNN, and here we use 10 neighbors) classifier. We split the gene set into training (80%) and testing (20%) datasets, and high accuracy (0.931) produced by the kNN classifier demonstrates that our embeddings are enriched with functional information. Extended Data Figures 4 and (b) display the functional clustering for these genes, where genes with similar functions are also embedded into a joint cluster. To further evaluate the correctness of functional clustering, we performed Gene Ontology Enrichment Analysis (GOEA) [47–49] and Ingenuity Pathway Analysis (IPA) [50] based on the top 10 clusters of protein-encoding genes ranked by the number of genes in clusters with the descending order. The top 1 cluster contains the largest number of genes. This setting can demonstrate the robustness of gene embeddings analysis based on different databases. We visualize the top pathways ranked by −*log*(adjusted p-value) (the p-value adjusted by Bonferroni correction) of each cluster in Extended Data Figures 5 and 9. We found that all the clusters contain pathways related to important biological activities, including metabolism, transport, and genetics, and others. We also analyzed text descriptions from NCBI and LLMs in Appendix A. The outputs of different LLMs are summarized in Supplementary File 1. We could also utilize embeddings from LLMs to analyze single-cell data and spatial transcriptomic data from Mouse and other species, thus offering potentials in cross-species analysis. These are summarized in Appendix B.

### scELMo for clustering and batch effect correction

In this section, we investigated the contribution of feature embeddings from scELMo for cell-level tasks including clustering and batch effect correction.

Clustering is effective in checking whether feature embeddings carry biological information. We incorporated gene embeddings or protein embeddings into the single-cell sequencing data and evaluated the performance of clustering based on cell embeddings. The metrics we chose include NMI, ARI, and ASW [52], which are widely used in the evaluation of clustering for single-cell data. In the following analysis, if not otherwise specified, GenePT means GenePT-w. Since we generated gene embeddings for gene names from both NCBI and Ensemble [53], we had more genes that those from GenePT. Moreover, when incorporating the feature embeddings, GenePT utilized a naive arithmetic average (*aa*) method, where it computed the embeddings for one cell by directly multiplying the gene expression vector from this cell and the matrix of gene embeddings from GPT 3.5. The *aa* mode ignored the scales of log-normalized gene expression levels across different cells. However, the level of gene expression is a key factor affecting cellular function [54, 55]. Different from GenePT, we treated the gene expression levels as weights and computed the weighted average (*wa*) value for each cell as an alternative averaging approach. We implemented these approaches for generating cell embeddings and details are included in the Methods section.

Using the *wa* mode was better than using the *aa* mode for cell clustering under different metrics, shown in Figure 2 (c). Meanwhile, the *wa* mode has better performances in three out of four datasets comparing with task-specific methods SC3 [56] and scVI [57]. Moreover, using embeddings from scELMo also improved the clustering performance compared with the embeddings from GenePT, advocating LLMs as a tool for summarizing scientific concepts. However, various approaches of combining the embeddings from GenePT and GPT 3.5 (e.g. GPT 3.5 + GenePT *wa* means sum by genes, and GPT 3.5 || GenePT *wa* means concatenation by genes.) did not improve the scores compared with the individual setting. The average ranks of all methods across different datasets are summarized in Extended Data Figure 6 (a), with the GPT 3.5 *wa* mode having the lowest rank. Finally, if we combined cell embeddings with the embeddings of cell-type information from GPT 3.5, we could get scores close to one, suggesting that the cell-type embeddings contained meaningful cell-type information. Appendix C summarizes factors that could affect clustering performance.

For the batch effect correction task, because GenePT already evaluated the performance of such embeddings for small-scale single-cell RNA sequencing (scRNA-seq) data, we first focused on single-cell proteomic data, such as those collected from CITE-seq [15] and CyTOF [14]. To ensure the fairness of benchmarking, our batch effect correction relies on projecting the datasets from different batches into a joint space without active training. Here we considered two scenarios. We tried to use the embeddings from scELMo and GenePT to reduce the batch effect for datasets from the same or different protocols. To evaluate the performance of batch effect correction, we used metrics from scIB [52] and considered the contribution of our embeddings for reducing batch effect (known as *S*_*batch*_) and for preserving the biological variation (known as *S*_*bio*_) separately. Details of our metrics are included in the Method section. We also included task-specific methods including MARIO [58], Harmony [59], and MNN [60] for comparisons. Figures 3 (a) and (b) display the results for batch effect correction from two CITE-seq datasets and Figures 3 (c) and (d) display the results for batch effect correction for one CITE-seq dataset and one CyTOF dataset. Based on these four figures, we find that using the *aa* approach did not improve the performance of scELMo for both cases. This observation also applied to task-specific methods. MARIO performed well in integrating the CITE-seq dataset and the CyTOF dataset, but it did not show superiority in integrating two CITE-seq datasets. MNN performed worse than either *aa* or *wa* approaches in these two datasets. Moreover, the *aa* approach could reduce *S*_*batch*_ and *S*_*bio*_ simultaneously compared with these two scores from the raw dataset under the Cytof+CITE-seq case, thus led to a larger batch effect. On the other hand, the *wa* approach led to the reduction of the batch effect as shown by the improvement of these two scores. Moreover, incorporating the cell-type embeddings with the *aa* mode also did not improve *S*_*batch*_ and *S*_*bio*_ jointly. However, combining the cell-type embeddings with the cell embeddings under the *wa* mode also improved the *S*_*bio*_ score for both cases. Therefore, finding a good base embedding space is important for utilizing cell-state embeddings. We also evaluated the performances of scELMo in integrating atlas-level scRNA-seq data [61–63], shown in Extended Data Figure 7. We demonstrated that scELMo outperformed GenePT in integrating large-scale transcriptomic data. As an extension, we summarize our analysis of multi-omic data integration [64] in Appendix D.

**Fig. 3.**
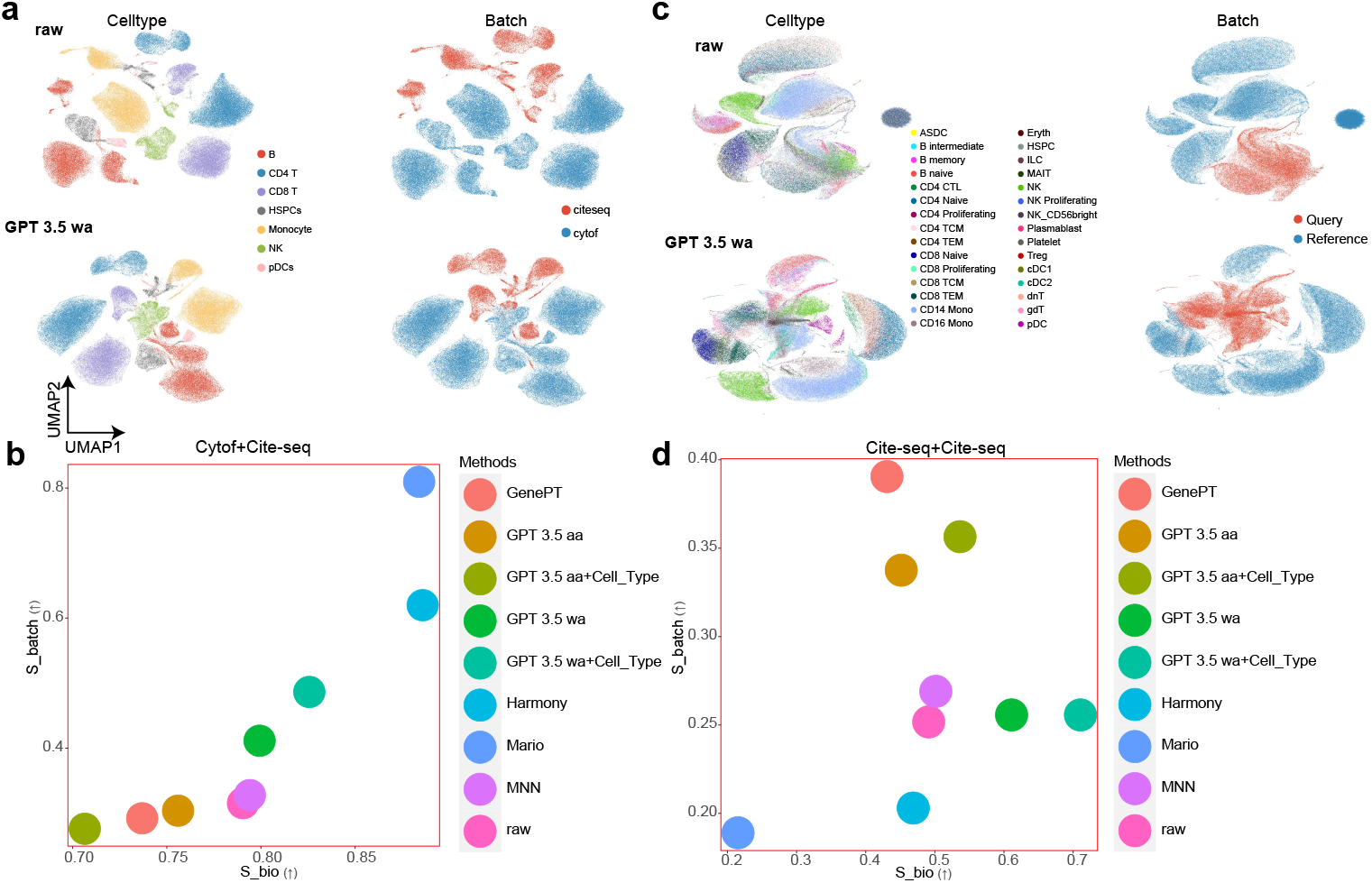
Results of batch effect correction for single-cell proteomic data. (a) UMAPs [51] for the cell-type information (left) and batch information (right) of cite-seq-based datasets. The upper panel represents the raw data. The bottom panel represents the cell embeddings from GPT 3.5 *wa* mode. (b) Evaluations of the batch effect correction for cite-seq-based datasets across different methods. (c) UMAPs for the cell-type information (left) and batch information (right) of cite-seq-based dataset and CyTOF-based dataset. The upper panel represents the raw data. The bottom panel represents the cell embeddings from GPT 3.5 *wa* mode. (d) Evaluations of the batch effect correction for cite-seq-based dataset and CyTOF-based dataset across different methods.

### scELMo for cell-type annotation

Cell-type annotation is a critical component of single-cell analysis [65]. Here we investigated the ability of scELMo to annotate cells under both the zero-shot learning framework and the fine-tuning framework. For the zero-shot learning framework, we incorporated the gene embeddings into both the training and testing datasets to access cell embeddings and used a kNN classifier implemented by GenePT to annotate cells in the testing dataset. However, this approach failed when the training datasets were from multiple resources or had large batch effects (shown in the PBMC section of Table 1). Therefore, we constructed a simple classifier based on neural networks [66] and contrastive learning [67], shown in Extended Data Figure 8 with description and explanation of model design. We then trained the model based on gene embeddings and expression profiles. Such model is also known as an adaptor in Natural Language Processing (NLP) tasks [29]. We tested the ability of cell embeddings from this model for annotating cell types. Here we considered three different datasets from different tissues for evaluation. The hPancreas dataset and the PBMC dataset contain one batch for testing and the rest of batches for training. The Aorta dataset is split into training and testing datasets from one study with a ratio of 0.8/0.2, by following the default setting of [3].

**Table 1.** Scores of cell-type annotation task under different settings. Parts of the results are directly extracted from GenePT. Here PCA represents principal component analysis, and *scELMo+random emb* represents fine-tuning scELMo with random numbers as meaningless gene embeddings. Average ranks of all methods across datasets are summarized in Extended Data Figure 6 (b). We boldfaced the highest score of each metric for each dataset.

|  | zero-shot settings | Accuracy (↑) | Precision (↑) | Recall (↑) | F1 |
| --- | --- | --- | --- | --- | --- |
| hPancreas [68-71] | GPT 2 query | 0.000 | 0.000 | 0.000 | 0.000 |
|  | GPT 4 query | 0.000 | 0.000 | 0.000 | 0.000 |
|  | scGPT (z) | 0.770 | 0.610 | 0.560 | 0.550 |
|  | Geneformer (z) | 0.500 | 0.250 | 0.340 | 0.270 |
|  | GenePT-w | 0.940 | 0.720 | 0.630 | 0.650 |
|  | GenePT-s | 0.890 | 0.650 | 0.530 | 0.560 |
|  | GPTCelltype | 0.501 | 0.386 | 0.415 | 0.366 |
|  | PCA | 0.633 | 0.369 | 0.382 | 0.357 |
|  | GPT3.5 aa | 0.933 | 0.702 | 0.614 | 0.629 |
|  | GPT3.5 wa | 0.933 | 0.702 | 0.613 | 0.629 |
|  | fine-tuning settings |  |  |  |  |
|  | scGPT | **0.970** | 0.742 | **0.758** | **0.741** |
|  | Geneformer | 0.321 | 0.105 | 0.094 | 0.086 |
|  | scELMo+GenePT | 0.963 | **0.790** | 0.662 | 0.680 |
|  | scELMo+random emb | 0.968 | 0.683 | 0.693 | 0.680 |
|  | scELMo+GPT 3.5 | **0.970** | 0.769 | 0.674 | 0.687 |
| Aorta [72] | zero-shot settings | Accuracy | Precision | Recall | F1 |
|  | GPT 2 query | 0.000 | 0.000 | 0.000 | 0.000 |
|  | GPT 4 query | 0.000 | 0.000 | 0.000 | 0.000 |
|  | scGPT (z) | 0.938 | 0.921 | 0.894 | 0.901 |
|  | Geneformer (z) | 0.860 | 0.700 | 0.600 | 0.620 |
|  | GenePT-w | 0.870 | 0.910 | 0.680 | 0.720 |
|  | GenePT-s | 0.860 | 0.700 | 0.600 | 0.620 |
|  | GPTCelltype | 0.340 | 0.254 | 0.261 | 0.228 |
|  | PCA | 0.929 | 0.923 | 0.911 | 0.910 |
|  | GPT3.5 aa | 0.877 | 0.867 | 0.678 | 0.719 |
|  | GPT3.5 wa | 0.877 | 0.867 | 0.677 | 0.718 |
|  | fine-tuning settings |  |  |  |  |
|  | scGPT | **0.962** | 0.938 | 0.939 | 0.937 |
|  | Geneformer | 0.340 | 0.087 | 0.092 | 0.080 |
|  | scELMo+GenePT | 0.958 | **0.952** | **0.940** | **0.946** |
|  | scELMo+random emb | 0.951 | 0.913 | 0.874 | 0.882 |
|  | scELMo+GPT 3.5 | 0.957 | 0.950 | 0.936 | 0.942 |
| PBMC [73-76] | zero-shot settings | Accuracy | Precision | Recall | F1 |
|  | GPT 2 query | 0.000 | 0.000 | 0.000 | 0.000 |
|  | GPT 4 query | 0.000 | 0.000 | 0.000 | 0.000 |
|  | scGPT (z) | 0.915 | 0.781 | 0.803 | 0.789 |
|  | Geneformer (z) | 0.126 | 0.128 | 0.130 | 0.095 |
|  | GenePT-w | 0.286 | 0.607 | 0.270 | 0.315 |
|  | GenePT-s | - | - | - | - |
|  | GPTCelltype | 0.501 | 0.387 | 0.415 | 0.366 |
|  | PCA | 0.436 | 0.392 | 0.278 | 0.285 |
|  | GPT3.5 aa | 0.190 | 0.564 | 0.181 | 0.230 |
|  | GPT3.5 wa | 0.190 | 0.562 | 0.181 | 0.231 |
|  | fine-tuning settings |  |  |  |  |
|  | scGPT | 0.933 | 0.798 | 0.822 | 0.807 |
|  | Geneformer | 0.235 | 0.146 | 0.147 | 0.131 |
|  | scELMo+GenePT | 0.919 | 0.785 | 0.823 | 0.801 |
|  | scELMo+random emb | 0.903 | 0.854 | 0.826 | 0.835 |
|  | scELMo+GPT 3.5 | **0.955** | **0.940** | **0.929** | **0.934** |

Table 1 shows that the zero-shot learning ability of embeddings from GPT 3.5 is impressive for annotating cells for the hPancreas dataset and the PBMC dataset. scELMo also outperformed GPTCelltype [4], which is based on the markers extracted from LLMs for cell-type annotation. We believe that GPTCelltype is limited by preclustering and pseudo-label assessment before calling the LLMs for annotation, which reduces the quality of marker genes, especially for complex datasets. However, scELMo argues that we can provide a better cell representation by integrating embedding information from LLMs, and we can improve cell-level tasks by using a better representation, and thus having an annotation set with higher quality. Moreover, representing cells by using the gene rank and the representation for GPT 2 or GPT 4 did not perform well. Therefore, zero-shot learning based on gene embeddings is capable of cell-type annotation for datasets with minimal batch effect. However, for the PBMC dataset, which consisted of datasets from different sources, there was a clear gap between zero-shot learning results and fine-tuning results. Moreover, the fine-tuning results by combining the gene embeddings from GPT 3.5 or GenePT and our adaptor are comparable with FMs like scGPT [1] and Geneformer [2]. However, scGPT and Geneformer need more resources including NVIDIA A100 GPU and longer running time for fine-tuning [18], further shown in Extended Data Figures 10 (a) and (b). For all the datasets we tested, scELMo with a fine-tuning framework performed better than the same settings with the exception of the zero-shot learning framework. Moreover, scELMo could improve the annotation score for embeddings from different sources. Our results show that cell-type annotation based on fine-tuning the adaptor with gene embeddings from LLMs is accurate and easy to implement.

### scELMo for in-silico treatment analysis

Using computational methods to discover novel target therapies and drugs has attracted much attention [77–79]. In this section, we study the ability of scELMo to use the adaptor for cell-type annotation and the gene embeddings we extracted from GPT 3.5 or NCBI to model human diseases and reveal potential therapeutic targets. This is a new task and has not been discussed in GenePT. Inspired by Geneformer [2], we finetuned our adaptor based on the classification task of cell conditions and used the cell embeddings from the adaptor for therapeutic target discovery. We utilized the training-validating framework to choose the adaptor with best performance and tested the change of embeddings by removing the differenetially expressed genes (DEGs) in the expression levels across different conditions. We discussed the importance of fine-tuning a model in Appendix E. Meanwhile we found that using gene embeddings from GPT 3.5 can give us the most accurate result in Figure 4(c). If we remove a suitable candidate for targeted therapies, the cell embeddings under disease will be more similar to those under control. The metric we used to evaluate similarity is the cosine similarity between the embeddings under the diseased case and the average cell embedding under the control case. We evaluated the discovered targets through GOEA and literature review as summarized in Supplementary File 2.

**Fig. 4.**
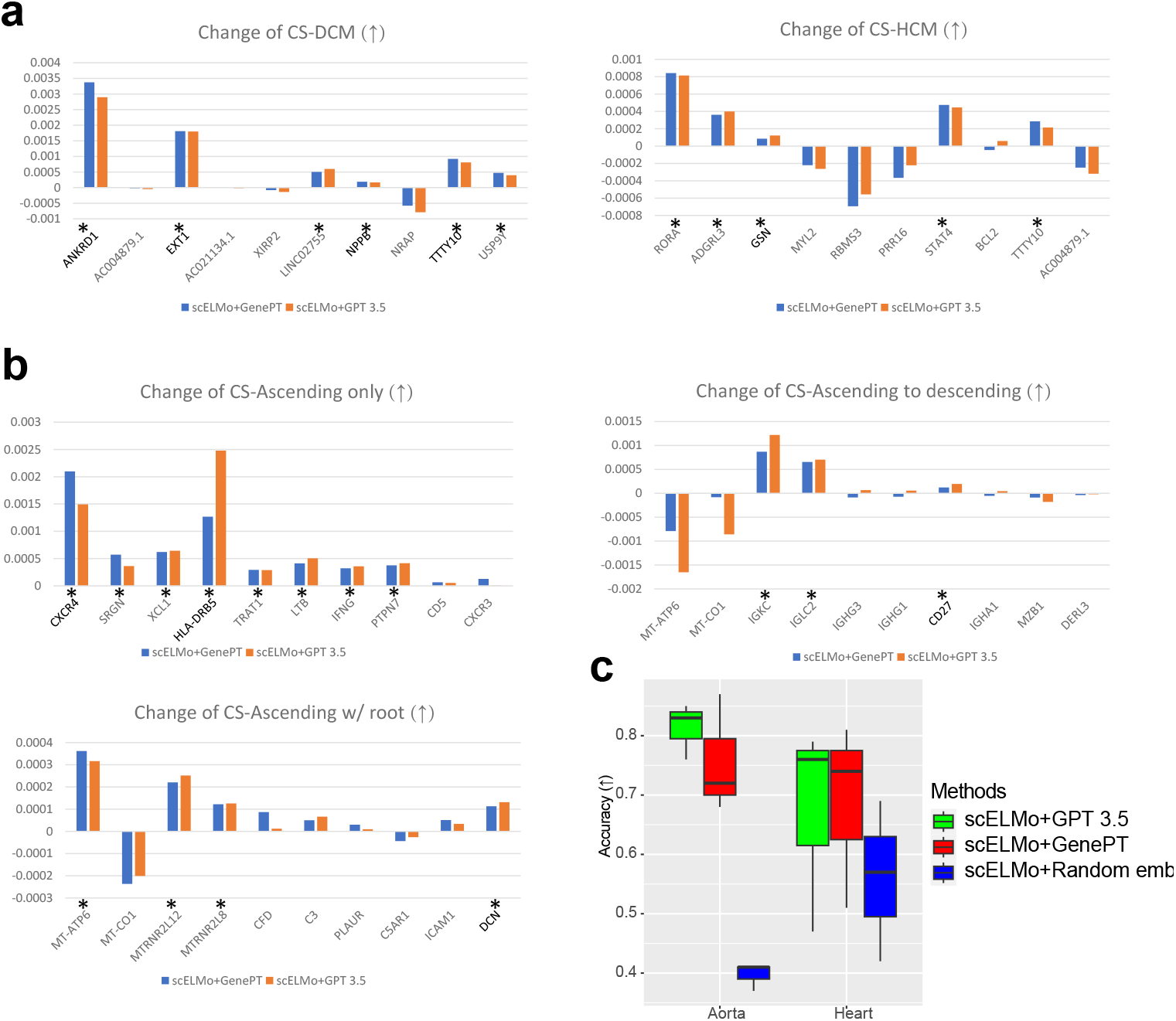
Results of in-silico treatment analysis. A gene is treated as a potential therapeutic target if the change of cosine similarity (CS) by removing the gene is larger than 1e-4. The reason for setting such threshold was discussed in the Methods section. We chose the top 10 DEGs as candidates. (a) The change of CS by removing different genes in the expression space for hypertrophic (HCM) or dilated cardiomyopathy (DCM) states. (b) The change of CS by removing different genes in the expression space for ascending aortic aneurysm (Ascending only, Ascending to descending, and Ascending w/root). (c) The accuracy of disease-state annotation under different settings of scELMo. We highlighted the genes detected by both GenePT and scELMo using stars (*) and marked the genes that were discovered by previous research as therapeutic targets using **bold** type.

We first considered hypertrophic or dilated cardiomyopathy states [80] using the scRNA-seq Heart dataset [81]. We identified genes whose in-silico deletion in the disease conditions could significantly shift the cell embeddings from the disease conditions to the non-failing (NF) or control condition. The change of cosine similarity before and after the deletion for cardiomyopathy states is summarized in Figure 4 (a). Our adaptor identified two novel potential therapeutic targets for cells under the DCM case and four novel targets for cells under the HCM case, and using gene embeddings from GPT 3.5 or NCBI gives us consistent result, shown in Figure 4 (a). The silence of genes identified by scELMo may help the shift from cells under the disease conditions to cells under the control condition. There is literature to support our discovery of ANKRD1 [82], EXT1 [83], NPPB [84] and TTTY10 [85]. Moreover, there is supporting evidence of GSN [2] by CRISPR-based technology [86] as a therapeutic target. We show the results of GOEA based on selected genes in Extended Data Figures 11 (a) and (b) with at least 10 pathways ranked by −*log*(adjusted p-value) and the results of IPA in Extended Data Figure 12 (a). According to the pathway information, the selected genes play an important role in cardiac activities, corresponding to the target tissue. Meanwhile, the select genes can also capture the interaction of these important pathways. These findings suggest the potential usefulness of genes selected by scELMo.

Then we considered ascending aortic aneurysm [87], which has three different disease states. The scRNA-seq dataset we used is the scRNA-seq Aorta dataset. We used the similar approach discussed above and summarize the change of cosine similarity before and after deletion in Figure 4 (b). scELMo identified six novel genes as potential therapeutic targets for cells under the Ascending only state, two novel genes as potential therapeutic targets for cells under the Ascending to descending state, and three genes as potential therapeutic targets for cells under the Ascending w/root state, shown in Figure 4 (b). Interestingly, MT-ATP6 gene was selected by scELMo as a potential target, which implied that cells under different states might have different speeds for cell death [88] or neurodegeneration [89]. We also show the results of GOEA based on selected genes in Extended Data Figure 11 (c)-(e) and the results of IPA in Extended Data Figure 12 (b). Our selected genes are important for immune responses according to these pathways. These results suggest the potential benefit of exploring the correlation between cellular function and clinical manifestations of this disease.

### scELMo for perturbation analysis

Analyzing the perturbation effect on cell state using single-cell data is also an important task, and since gene embeddings from LLMs contain functional information of possible perturbed targets, we intend to explore the contributions of scELMo for perturbation modelling and analysis. Here we focus on three tasks across different perturbation types, for example, chemical perturbation [93] or gene-level perturbation (perturb-seq) [94]. Our idea is to use the cell embeddings or gene embeddings generated based on the zero-shot learning framework of scELMo to replace or combine with the original input of different models to enhance their performance on the tasks including causal factor analysis, gene expression prediction with chemical perturbation and gene expression prediction with gene-level perturbation. We evaluated the contribution of scELMo for the causal factor analysis task based on CINEMA-OT [92], the gene expression prediction task for chemical perturbation data based on CPA [95], and the gene expression prediction task for gene-level perturbation data based on GEARS [96]. All of the metrics we considered in these three tasks are from these models.

CINEMA-OT is a causal learning framework based on optimal transport that separates perturbation effects from intrinsic cell-state effects in scRNA-seq datasets. To evaluate the contribution of our embeddings, we replaced CINEMA-OT’s default input (PCA features) with cell embeddings from scELMo. First, in the intrinsic cellstate (confounder) space where perturbation effects are removed, embeddings from the same cell type should not differ by perturbation cases. Using our cell embeddings as input improved CINEMA-OT’s performance compared with PCA, leading to reduced batch effects for cells of the same type, shown in Figure 5 (a). Furthermore, in this setting, GenePT gene embeddings outperformed gene embeddings derived from GPT-3.5. Second, we examined whether gene embeddings alone are sufficient for causal factor analysis across different cell types. As shown in Figure 5 (b), relying solely on gene embeddings did not significantly improve the separation of perturbation effects and intrinsic cell states. In contrast, when we integrated both gene-level and cell-type embeddings, the modified CINEMA-OT achieved substantially better separation across perturbation cases. This improvement was supported by Wilcoxon Rank-Sum tests comparing GenePT (default) and scELMo embeddings, yielding significant differences in two datasets (p = 0.031 for both cases).

**Fig. 5.**
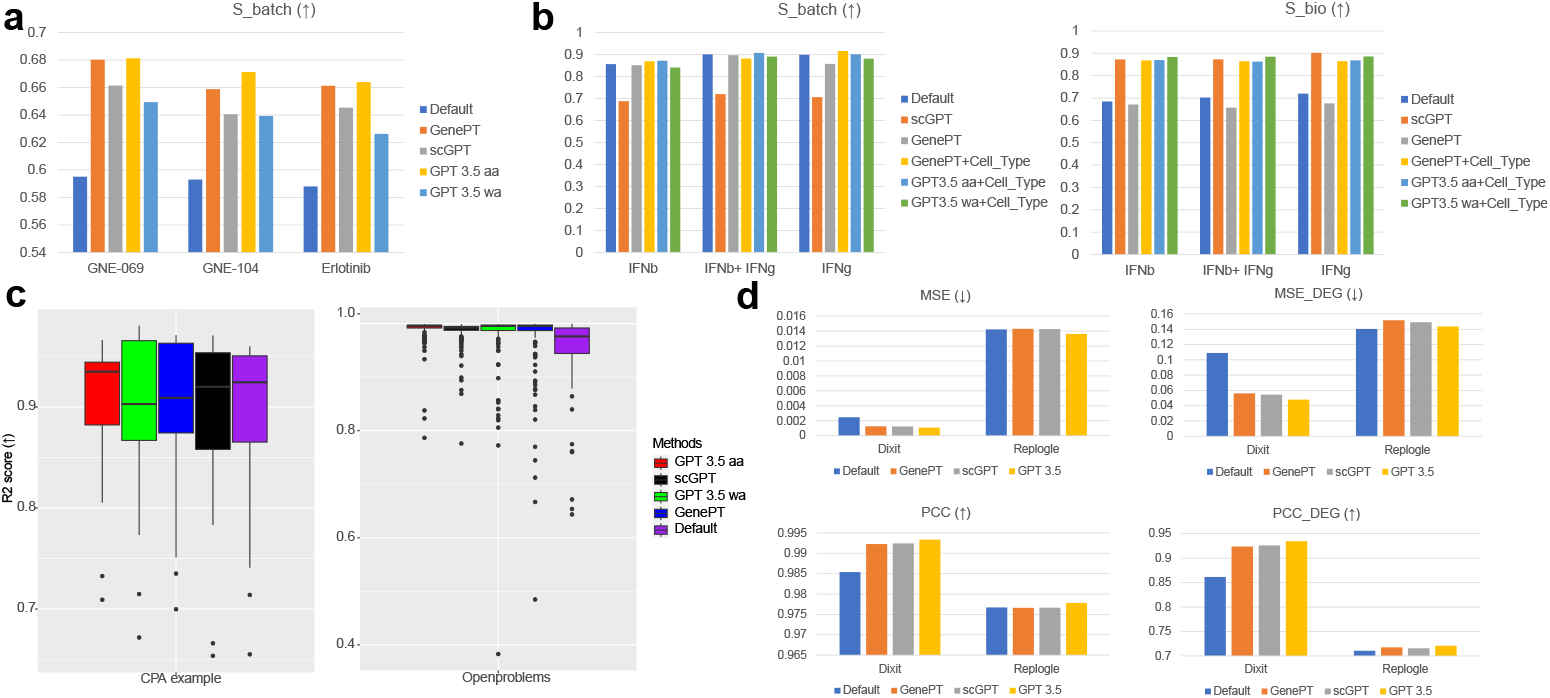
Results of perturbation analysis. (a) Scores of casual factor analysis for ChangYe2021 dataset [90, 91] based on CINEMA-OT across different input data. We considered four different types of extension. We still use *S*_*batch*_ to represent the levels of perturbation effect removal because we used the metrics for benchmarking batch integration. (b) Scores of causal factor analysis for perturbed PBMC dataset [92] based on CINEMA-OT across different input data. (c) Scores of gene expression prediction under different perturbation cases based on CPA. We considered four different methods and two datasets. (d) Scores of gene expression prediction using perturb-seq datasets based on GEARS. We considered three different methods and three datasets. The arrows represent the direction of metrics comparison.

Moreover, Extended Data Figures 13 (a) and (b) show that the visualization could also be improved by using the updated embeddings as model input. Extended Data Figures 13 (c) and (d) show that incorporating the cell embeddings into CINEMA-OT did not affect the analysis of gene synergy effect, and the difference between synergy of Monocyte and synergy of other cell types was also obvious. For the causal factor analysis task, using the *wa* mode did not improve the score. One possible reason is that CINEMA-OT can learn the best representation of cell embeddings with a good start for optimization, so the *aa* mode is adequate. Therefore, scELMo can improve the performance of CINEMA-OT on the causal factor analysis task by offering a new candidate of input data.

CPA is a tool based on Conditional Variational Auto-encoder (CVAE) to predict the gene expression levels for the out-of-distribution (OOD) samples of scRNA-seq data under chemical perturbations. Here we combined the gene embeddings from scELMo with the original input dataset and learned a new latent space for gene expression prediction. We investigated the contribution of gene embeddings by comparing the R2 score between these two different settings. We computed the R2 score based on the predicted gene expression levels and the observed gene expression levels. Figure 5 (c) shows the performance of CPA under different methods for two datasets. For the CPA example dataset shown in the left panel, using the cell embeddings from GenePT and the GPT 3.5 in the training process slightly improved the average R2 score, while its median value was still lower than the default mode. For the Openproblems dataset [97] shown in the right panel, using cell embeddings from both scGPT, GPT 3.5 and GenePT improved the performance of CPA. Moreover, the R2 score based on combining cell embeddings with CPA had a higher average value and lower variance compared with the default mode. We further performed Wilcoxon Rank-sum tests for the R2 scores between different conditions and set the significant threshold for p-value as 0.05. Based on our computation, the pairs of settings with significant differences include GPT 3.5 aa vs. Default (p-value=0.003), GPT 3.5 aa vs. GenePT (p-value=0.002) based on the CPA example dataset, and GPT 3.5 aa vs. Default (p-value=7e-11), GPT 3.5 wa vs. Default (p-value=6e-12) based on the Openproblems dataset. Therefore, scELMo could improve the performance of CPA on the prediction task by introducing the cell embeddings into the training process.

GEARS is a tool based on Graph Neural Networks (GNN) [98] to predict the gene expression levels for perturb-seq-based datasets. Here we combined the gene embeddings from scELMo with the original gene embeddings of GEARS to learn the predicted value of target genes. We studied the contribution of gene embeddings by comparing the Pearson Correlation Coefficient (PCC) and Mean Squared Errors (MSE) between the default settings and updated settings when all genes were considered and when only DEGs were considered. The results are summarized in Figure 5 (d) and Extended Data Figure 14 (a). For both Replogle [99] and Dixit [94] datasets, scELMo based on gene embeddings from GPT 3.5 outperformed the default settings as well as scELMo based on gene embeddings from GenePT and scGPT, supported by the comparisons based on all the metrics. For the Dixit dataset, using gene embeddings from GPT 3.5 improved the gene expression prediction made by GEARS for both the all genes case and the DEGs case obviously. By considering the four datasets (Dixit, Norman [100], Adamson [101], and Replogle [99]) together, we performed Wilcoxon Rank-sum test between the PCC and PCC DEG of scELMo and the default mode, and the improvement of scELMo is significant (p-value=0.027). Furthermore, we also simulated datasets with different numbers of cells based on the Dixit dataset to test the robustness and consistency of our improvement, inspired by [102]. Extended Data Figure 14 (b) shows that scELMo has an overall better performance as well as a lower variance compared with the default mode. Therefore, we believe that the introduction of gene embeddings from LLMs can also contribute to perturbation effect prediction by incorporating more knowledge about perturbed genes.

## 3 Discussion

Developing methods to model genetic and cellular functionalities is an important task in computational biology. Although a common practice is to propose a biological question and design relevant experiments to answer this question, recent years have seen the development of a foundation model that may be informative to address many known and unknown biological questions. In the single-cell data analysis area, many researchers have proposed to pre-train a large-scale model and declare it as a foundation model by showing their comparable performance with different task-specific models in specific downstream tasks. However, it is hard to find tasks that could only be resolved through such pre-training and fine-tuning framework [18], which questioned the contribution of consuming large-scale resources to develop such models. Therefore, inspired by GenePT, we explored another approach to generalizing a foundation model in the single-cell research area, that is, utilizing the contribution of LLMs to generate meaningful feature embeddings and cell embeddings. We could either directly utilize these embeddings for clustering or reducing batch effect, or combine these embeddings with task-specific models to improve their performance. These two approaches are integrated in scELMo. Our gene embeddings are also robust to randomness, shown in Extended Data Figure 15 for different downstream tasks with low variance. Accessing the outputs of known LLMs like GPT 3.5 did not require many resources, and our results did show the superiority of scELMo in answering multiple biological questions.

For scELMo under the zero-shot learning framework, we could utilize cell embeddings to perform clustering and batch effect correction. These contributions are based on the fact that embeddings from the text description of features in single-cell datasets are good for representing biological concepts or functions. We also discussed the factors that could affect the performance of generating meaningful cell embeddings, including the approach to computing the average embeddings, the number of cells in one dataset, the number of features we need, and other factors. We showed that such embeddings could be used for multi-omic data analysis, which illustrates the power of using LLMs as tools for incorporating prior information to enhance task-specific analysis.

Considering the limitation of the zero-shot learning framework, we also proposed a fine-tuning framework for scELMo. By combining the feature embeddings from GPT 3.5 with a light-structured neural network, we could use the embeddings to annotate cell types with performance similar to those of FMs that require much more resources for pre-training and fine-tuning. Moreover, scELMo can also be used for detecting novel therapeutic targets by examining the change of embeddings corresponding to the removal of certain genes, supported by related biological experiments. We could also directly incorporate the cell embeddings or feature embeddings with task-specific models for better performance in modeling the data with perturbation. In-silico treatment analysis and perturbation analysis are two challenging and important tasks with cell-level knowledge, which further supported the potential of scELMo and related work.

Here we suggest a guideline for users interested in scELMo. The *wa* mode works better for clustering and batch effect correction, while the *aa* mode works better for perturbation analysis. We explain the rationale of such setting in the Method section. Both types of embeddings can help in performing cell-type annotation and insilico treatment analysis. We recommend users consider the embeddings of meta-data information based on their specific tasks and requirements. We also encourage users to actively explore the boundaries of scELMo’s capabilities to discover more interesting applications towards biology.

However, scELMo also has the following limitations. Firstly, the rapid developments of LLMs likely lead to embeddings better than GPT 3.5. With a more powerful LLM, we will have a better representation of features and cells. Secondly, LLMs could not generate meaningful information for genes that were recently discovered or analyzed. Although GPT 3.5 does not make up concepts for genes, the lack of knowledge still presents a question for the applications of scELMo. This shortcoming might limit the performance of applying scELMo for low-resource biological data, such as cells from patients with rare diseases. Finally, extracting features from other biomedical data like GWAS [103] or scATAC-seq [104] data will be difficult since the number of features for these data is quite large.

Therefore, in the future, we plan to generate a database containing the text description as well as the embeddings of such descriptions of features from different LLMs. Models like CPA can utilize these embeddings to perform prediction under the OOD cases. Considering the limitation of our embeddings to enhance the perturbation prediction task, we may incorporate the perturbation information of target genes’ descriptions or consider better metrics for evaluating tasks related to perturbations. Moreover, we will keep on figuring out a more practical approach to offer a gene-specific LLM. In addition, since the idea from scELMo is capable of modeling arbitrary biomedical data with tabular format, we believe that extending the usage of scELMo to other tasks or areas is also very promising.

## 4 Methods

### 4.1 Problem definition

For a typical single-cell dataset *X*^*n×m*^ after normalization [105] with *n* cells and *m* features, our target is to utilize the text description from a mapping function ℳ () for feature-level metadata information *f*^*m×*1^ and cell-level metadata information *c*^*n×*1^ to learn the embeddings of cells. Each cell or gene has one corresponding metadata information description and thus *f* and *g* are vectors. If we define the embedding generation layer of ℳ () as ℳ _e_(), our cell embeddings (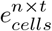, where *t* represents the dimension of LLM embeddings) can be represented as:

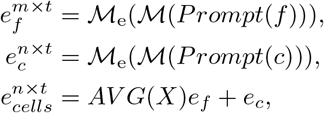

where *Prompt* is a mapping function that can transfer the name of input data to the prompt space. The prompts can be used as the input of language models. The function *AV G*() represents the method we used to average the embeddings of all genes for each cell. If the mode is *aa*, we divide *X* by *m*. If the mode is *wa*, we divide each row of *X* by the sum of this row. Considering the cell with index *i*, we can define these two processes as:

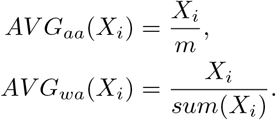

Then we use matrix multiplication to combine the feature embeddings and the expression profile. Our default setting of the mapping function is a LLM. GenePT can be treated as a special case of scELMo, that is, replacing the LLM with a known database and using *aa* mode. Incorporating the embeddings of cell-level metadata is an optional choice. We intend to investigate if the cell embeddings can offer a better representation than the raw data.

Moreover, with the embeddings of feature-level metadata information and cell-level metadata information, we also consider if incorporating our embeddings with task-specific model 𝒯 can improve the performance of 𝒯, that is:

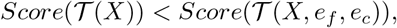

where *Score*() is a metric to evaluate the output of the given model, where higher value represents better output. We also may not need to have *e*_*f*_ and *e*_*c*_ for every model.

### 4.2 Method explanation

The default embedding model to generate embeddings of text descriptions is from OpenAI as a closed-source model. Our consideration of *AV G* is inspired by [3] as well as the noise level of scRNA-seq dataset [106]. By compressing the gene expression profiles into a lower dimension, we can jointly model both the magnitude of gene expression levels and the functional information of genes. Moreover, we assume and demonstrate that such new cell embeddings can represent cells in a better approach contributed by the external information of gene functions and cell-type-specific information generated by LLMs. The raw data, on the other hand, are no longer required for further analysis. Furthermore, a systematical evaluation of LLM embeddings for medical machine learning usage [107] also supports our assumption. We found that *wa* mode worked better in handling the tasks of clustering and batch effect correction. As a possible reason, we believe that we need to consider the contributions of individual genes as well as their grouped effect for generating the cell embeddings under the zero-shot learning framework. For the fine-tuning framework, different task-specific methods have their favorable designs to incorporate LLM embeddings as prior knowledge, and thus we have a more flexible design.

### 4.3 scELMo under the zero-shot learning framework

To evaluate the performance of *e*_*cells*_ under the zero-shot learning framework, we consider three tasks: clustering, batch effect correction, and cell-type annotation based on a kNN classifier. In this section, 𝒯 is defined as a kNN classifier.

For clustering and batch effect correction, we directly use *e*_*cells*_ as a new representation for *X* and evaluate *e*_*cells*_ for these two tasks. For cell-type annotation based on a kNN classifier, we consider training dataset *X*_*train*_ and testing dataset *X*_*test*_, and their corresponding cell embeddings 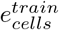 and 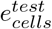. We use 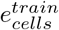 with its cell types to train a kNN classifier and perform cell-type annotation based on 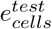. Since kNN is based on similarity searching, we treat this method as an ability of zero-shot learning. We follow the settings from GenePT for this classifier and set k=10.

### 4.4 scELMo under the fine-tuning framework

To evaluate the performance of *e*_*f*_ and *e*_*c*_ under the fine-tuning framework, we considered three tasks: cell-type annotation with an adaptor, in-silico treatment analysis with an adaptor, and perturbation analysis with task-specific models as adaptors.

For the cell-type annotation and in-silico treatment analysis tasks, we propose a light-structured neural network with a contrastive learning [67, 108] design. Here 𝒯 is a neural network with ReLU [109] as the activation function. Our intuition comes from the requirement for a good representation of cells with different labels and conditions. Therefore, we formalize the loss function of our model as:

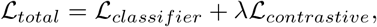

where ℒ_*classifier*_ represents the classification loss of the model output as we use cell-type labels for model training, and ℒ_*contrastive*_ represents the contrastive learning loss we use to distinguish the representations of cells under different conditions in the latent space. *λ* is a hyper-parameter, where we set *λ* = 100 in this manuscript to assign a larger weight for label-aware clustering. We utilize the embeddings after fine-tuning as the training and testing datasets and use a kNN classifier to annotate the cell types to evaluate the representation we learn based on scELMo.

To analyze the target of in-silico treatment, we first compute the cosine similarity (*CS*_*old*_) between the average cell embeddings based on the diseased case and control case. Then we delete the target gene by setting its expression profile as zero and compute the new embeddings and cosine similarity (*CS*_*new*_). We define the score of our targeted gene *g* as:

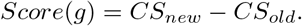

If such a score is larger than 1e-4, we treat the gene we analyze as a candidate for therapeutic targets. This threshold is based on the upper bound of the tiny quantities determined by Numpy [110] for scientific notation representation and the smallest non-zero scale of the y-axis in Figures 4 (a) and (b). We utilize the selected genes to run gene pathway analysis based on Gene Ontology Enrichment Analysis (GOEA) [47–49] and Ingenuity Pathway Analysis (IPA) [50].

For the perturbation analysis task, we consider three different models for the three tasks. Here 𝒯 represents different models corresponding to different perturbation analysis tasks. For the causal factor analysis task and CINEMA-OT, we replace the original input of CINEMA-OT with *e*_*cells*_. We follow the default settings of CINEMA-OT for processing related datasets. For the gene expression prediction task and CPA, we add a new neural network component to make *e*_*cells*_ learnable and combine the output of this component with the latent space from the original CPA. We do not modify the training process of CPA. We follow the default settings of CPA for processing related datasets. For the gene expression prediction task based on perturb-seq-based datasets and GEARS, we add the *e*_*f*_ to the original gene embeddings of GEARS. We do not modify the training process of GEARS. We follow the default settings of GEARS for processing Dixit, Norman, Adamson, and Replogle datasets.

### 4.5 Data pre-processing

We follow the data pre-processing steps from Scanpy [105] for scRNA-seq datasets. For single-cell proteomic datasets, we follow the pre-processing steps from TotalVI [111] and MARIO [58] and do not change the distribution of the original data because of its density.

#### Metrics

To evaluate the hallucinations of LLM outputs, we consider two metrics including BiLingual Evaluation Understudy (BLEU) [43] score and Human-Eval score [44].

The BLEU score is used to evaluate the similarity between observed string *ŷ* and ground truth string *y* based on *n* − *grams* strategy. Considering we have a function *C*(*s, y*) to generate the number of appearances of *s* as a substring of *y* and a set *G*_*n*_(*ŷ*) as the *n* − *grams* set, the BLEU score is defined as:

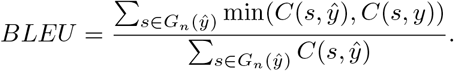

The score is in [0,1] and a higher value means better performance.

The Human-Eval score means we compare the truthfulness between the LLM outputs and references from NCBI and GeneCard databases to assign scores for the string pairs. We assign 1 if the outputs contain the correct information and 0 if the outputs do not contain the correct information. We have one human expert for evaluation. A higher value means better performance.

For the evaluations of clustering and batch effect correction, we utilize the metrics described and implemented by scIB [52]. We compute all the metrics we could compute in the evaluation process. All of the scores of the metrics in scIB are in [0,1] and a higher value means better performance.

For clustering, we use NMI, ARI, and ASW_*label*_ for evaluation. For batch effect correction, we compute ASW_*batch*_, PCR, Graph Connectivity, kBET, and iLISI and average the scores from these metrics to generate *S*_*batch*_. We compute ASW_*label*_, NMI, ARI, and cLISI and average the scores from these metrics to generate *S*_*bio*_. Details of these metrics, referred from [18, 52], are introduced below:

##### 1. Normalized Mutual Information (NMI)

NMI is a score to evaluate the performance of biological information conservation. We compute this score based on the mutual information between the optimal Leiden clusters and the known cell-type labels and then take the normalization. NMI ∈ (0, 1) and higher NMI means better performance.

##### 2. Adjusted Rand Index (ARI)

ARI is a score to evaluate the performance of biological information conservation. ARI is used to evaluate the agreement between optimal Louvain clusters and cell-type labels. ARI ∈ (0, 1) and higher ARI means better performance.

##### 3. Average Silhouette Width (ASW)

We have cell type ASW (*ASW*_*cell*_) and batch ASW (*ASW*_*batch*_) for this metric. For one cell, ASW calculates the ratio between the inner cluster distance and the intra cluster distance for this cell. Therefore, higher *ASW*_*cell*_ means better biological information conservation and lower *ASW*_*batch*_ means better batch effect correction. To make them consistent, for *ASW*_*cell*_, we take the normalization, that is:

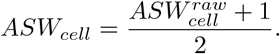

Similarly, for *ASW*_*batch*_, we take the inverse value of the normalized result, that is:

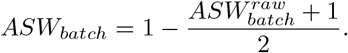

Both of metrics are in (0,1), and a higher score means better model performance.

##### 4. Local Inverse Simpson’s Index (LISI)

LISI is a metric to evaluate whether datasets are well-mixed under batch labels (*iLISI*) or can be discerned with different cell types *cLISI*. We first compute the k-nearest-neighbor list of one cell and count the the number of cells that can be extracted from the neighbors before one label is observed twice. Furthermore, we take the normalization for *iLISI* with *B* batches, that is:

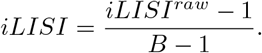

Similarly, for *cLISI* with *C* cell types, we take the inverse value of the normalized result, that is:

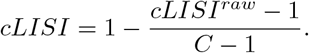

Both of metrics are in (0,1), and a higher score means better model performance.

##### 5. Graph Connectivity (GC)

GC measures the connectivity of cells in different cell types. If the batch effect is substantially removed, the connectivity of cells of the same cell type from different batches will have a higher connectivity score based on the k-NN neighbor graph. Therefore, we can compute the GC score for each cell type and take the average. GC score is in (0,1) and higher means better batch effect correction performance.

##### 6. Principal Component Regression (PCR)

PCR is a metric to evaluate the performance of batch effect correction. We calculate the *R*^2^ for a linear regression of the covariate of interest onto each principal component. The variance contribution of the batch effect for all the PCs is based on the sum of the product between the variance of each PC and the *R*^2^ of each PC across all the PCs. Therefore, the score can be represented as:

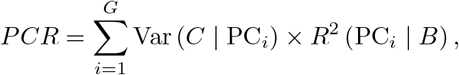

where *G* denotes the number of PCs and *B* denotes the batch information. PCR is in (0,1) and a higher score means better performance.

##### 7. kBET

the kBET algorithm is used to determine if the label composition of the k-nearest-neighbors of a cell is similar to the expected label composition. For the batch label mixture of cells in the same cell types, the proportion of cells from different batches for the neighbors of one cell should match the global level distribution. The kBET score ∈ (0, 1) and higher score means better batch effect correction performance. For the evaluations of cell-type annotation, we use Scikit-learn [112] to calculate Accuracy, Precision, Recall, and F1 score by comparing the predicted cell-type labels and ground-truth cell-type labels. All of the metrics are in [0,1] and a higher value means better performance.

For the evaluations of in-silico treatment analysis, we use Scipy [113] to compute the cosine similarity between the mean cell embeddings from the control case and the mean cell embeddings from the diseased case. The definition of the score here is described in the Methods section.

For the evaluations of perturbation analysis, we have three different tasks with different metrics. For the causal factor analysis task, the metrics are the same as what we use in the batch effect correction task. For the gene expression prediction task based on CPA, we use the R2 score as a metric to evaluate the performance for regression. R2 score is defined as:

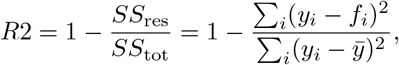

where *y*_*i*_ represents the ground-truth gene expression level, *f*_*i*_ represents the predicted gene expression level, and 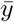 represents the average expression levels of the given gene. A higher averaged R2 score and a lower variance mean better performance.

For the evaluation of gene expression prediction tasks based on GEARS, we use PCC and MSE as metrics. We define PCC as:

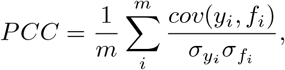

where *m* represents the number of used genes, *cov* represents the covariance, *σ* represents the standard deviation. *ρ* is in [0,1] and a higher value means better performance.

We can also define MSE as:

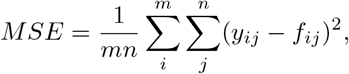

where *y*_*ij*_ represents the ground truth gene expression level of gene *i* in cell *j*, and *f*_*ij*_ represents the predicted gene expression level of gene *i* in cell *j*. Lower MSE score means better performance.

We consider computing these two metrics for both the all genes case and the top 20 DEGs case.

### 4.6 Explanations of baseline models

For tasks related to description generation, we consider MetaPrompt [45] and Chain-of-Thought (COT) [45] as baseline models for prompt engineering. MetaPrompt introduces a system prompt for LLMs and generates the outputs conditioned on the context in the system prompt. COT allows LLMs to obtain complex reasoning capabilities by allowing models to address the problem with intermediate steps.

For tasks related to clustering, we consider PCA (raw) [112], GenePT [3], SC3 [56], and scVI [57] as baseline models. PCA is widely used for dimension reduction of single-cell data. The principles of GenePT are summarized in the Introduction section. SC3 utilizes consensus clustering for analyzing scRNA-seq data, which is based on aggregating the k-means results after PCA transformation to generate the consensus and then performs the clustering based on consensus. scVI utilizes a generative model to learn the embeddings in the latent space of scRNA-seq data.

For tasks related to batch effect correction, we consider PCA (raw), GenePT, MARIO [58], Harmony [59] and MNN [60] as baseline models. MARIO considers both the shared features and distinct features for proteomic data and performs paired matching for cells from different batches to generate integrated results in the latent space. Harmony assigns labels for different cells with a soft-clustering method, computes the centroids for different clusters, and updates cell embeddings based on the soft cluster membership. MNN utilizes mutual nearest neighbors to learn the relationships for cells in different batches and updates the cell embeddings based on the relationships.

For tasks related to cell-type annotation, we consider GPT 2 [36], GPT 4 [23], scGPT [1], Geneformer [2], GPTCelltype [4], Multilayer Perception (MLP) and PCA for evaluation. To evaluate GPT 2 and GPT 4, we transfer the cell information into a sentence based on the rank of genes for each cell and treat this task as a Question-Answer task. scGPT is a pre-training-based FM for multiple tasks in single-cell research. The authors utilized multi-layer transformers to construct the model architecture and pre-trained scGPT based on large-scale scRNA-seq datasets. They finetuned scGPT to address problems in downstream applications like cell-type annotation. Geneformer is also a model based on pre-training. The authors utilized BERT [114] as a based model and pre-trained BERT from the sketch using scRNA-seq data transferred into sentences. They also finetuned Geneformer to address problems in downstream applications including cell-type annotation, in silico-treatment analysis, etc. GPTCelltype utilizes GPT-4 to extract markers of cell types to annotate cell clusters. To evaluate this model, we unify the model outputs and ground truth cell-type labels. MLP means we fits a MLP based on gene expression profiles and cell types. We extract the embeddings from these two models and PCs based on both zero-shot learning framework and fine-tuning framework, and utilize kNN to perform classification based on the embeddings.

For tasks related to perturbation analysis, we consider CINEMA-OT [92], CPA [95], and GEARS [96] for modification and evaluation. We included gene embeddings from GPT 3.5, NCBI (GenePT), and scGPT as inputs. CINEMA-OT is a method for separating confounding signals or causal factors from perturbations at single-cell level. The first step is to initialize the expected matrix rank based on biwhitening [115]. The second step is to separate confounder signals and treatment-associated signals based on Independent Component Analysis (ICA) [112]. The last step is to match the cells based on entropy regularized optimal transport [116]. We analyze the confounder embeddings for CINEMA-OT and follow the benchmarking pipeline mentioned in the original paper. CPA is a method for modeling the gene expression levels of scRNA-seq data with perturbations. CPA treats perturbations of cells as covariates and encodes these covariates as embeddings into the training process of a CVAE. CPA can also be used to predict the gene expression levels of OOD samples. GEARS is a method for predicting gene expression levels of perturb-seq-based datasets. It models the perturbations of genes based on knowledge graph and gene embeddings, thus utilizing GNN and Multi-layer Perceptrons (MLPs) to predict the gene expression levels under gene-level perturbation.

## 5 Codes and Datasets

The codes of scELMo can be found in https://github.com/HelloWorldLTY/scELMo. The license is MIT license. To generate the text descriptions and embeddings, we rely on the API of OpenAI. To run scELMo, we rely on Yale High-performance Computing Center (YCRC) and utilize one NVIDIA A5000 GPU with up to 30 GB RAM for fine-tuning. We utilize the bigmem node with up to 1000 GB RAM for analysis, and the minimal RAM requirement to analyze *∼*1,000,000 cells is 70 GB.

We did not generate new sequencing datasets in this project. All of the scRNA-seq data used in this study can be downloaded using the following links:

- https://github.com/JackieHanLab/TOSICA
- https://drive.google.com/drive/folders/1LgFvJqWNq9BqHbuxB2tYf62kXs9KqL4t
- https://academic.oup.com/bioinformatics/article/38/16/3942/6623406
- https://www.biorxiv.org/content/10.1101/2022.05.09.490241v2.abstract
- https://www.ncbi.nlm.nih.gov/geo/query/acc.cgi?acc=GSE139369
- https://www.ncbi.nlm.nih.gov/geo/query/acc.cgi?acc=GSE174072
- https://www.nature.com/articles/s41591-021-01329-2
- https://www.nature.com/articles/s41591-023-02327-2
- https://www.nature.com/articles/s41586-020-2797-4
- https://www.nature.com/articles/s41586-022-04817-8
- http://projects.sanderlab.org/scperturb/datavzrd/scPerturb_vzrd_v1/dataset_info/index_1.html
- https://github.com/vandijklab/CINEMA-OT/tree/main
- https://www.kaggle.com/competitions/open-problems-single-cell-perturbations
- https://cellxgene.cziscience.com/e/3faad104-2ab8-4434-816d-474d8d2641db.cxg/
- https://www.nature.com/articles/s41587-023-01905-6
- https://www.sciencedirect.com/science/article/pii/S0092867422005979?via#3Dihub

The download links and data statistics of all data are summarized in the Supplementary File 3. Source data of presented figures are available in Supplementary File 4.

## 6 Acknowledgments

We thank Gefei Wang and Chen Lin for the suggestions of datasets. We thank Mingze Dong for helpful discussions. This project is supported in part by NIH grants U24HG012108 and U01HG013840.

## 7 Author contributions

T.L. designed this study. T.L., X.L. and Y.C. designed the model. T.L., T.C. and W.Z. ran all the experiments. T.L. and H.Z. wrote the manuscript. H.Z. supervised this work. We did not use AI tools to write this manuscript.

