## Supplementary figures, supplementary files 1-4 for "scELMo: Embeddings from Language Models are Good Learners for Single-cell Data Analysis": Appendix.pdf

### A Examples of the text description outputs of different LLMs.

In this section, we present the differences between the text description from NCBI and the text description from GPT 3.5 for the same gene. We highlight the problematic output information of each text using red color, the description of functional information of each text using blue color, and the specific information contained in each text using **bold** text.

Here is an example for the text description of gene COL1A1 from NCBI:

*Official Symbol COL1A1provided by HGNC Official Full Name collagen type I alpha 1 chainprovided by HGNC Primary source HGNC:HGNC:2197 See related* ***Ensembl:ENSG00000108821 MIM:120150; AllianceGenome:HGNC:2197 Gene type protein coding RefSeq status REVIEWED Organism Homo sapiens Lineage Eukaryota; Metazoa; Chordata; Craniata; Vertebrata; Euteleostomi; Mammalia; Eutheria; Euarchontoglires; Primates; Haplorrhini; Catarrhini; Hominidae; Homo Also known as OI1; OI2; OI3; OI4; EDSC; CAFYD; EDSARTH1*** *Summary This gene encodes the pro-alpha1 chains of type I collagen whose triple helix comprises two alpha1 chains and one alpha2 chain. Type I is a fibril-forming collagen found in most connective tissues and is abundant in bone, cornea, dermis and tendon. Mutations in this gene are associated with osteogenesis imperfecta types I-IV, Ehlers-Danlos syndrome type VIIA, Ehlers-Danlos syndrome Classical type, Caffey Disease and idiopathic osteoporosis. Reciprocal translocations between chromosomes 17 and 22, where this gene and the gene for platelet-derived growth factor beta are located, are associated with a particular type of skin tumor called dermatofibrosarcoma protuberans, resulting from unregulated expression of the growth factor. Two transcripts, resulting from the use of alternate polyadenylation signals, have been identified for this gene. [provided by R. Dalgleish, Feb 2008] Expression Biased expression in gall bladder (RPKM 850.7), urinary bladder (RPKM 497.1) and 11 other tissues See more Orthologs mouse all.*

Here is an example for the text description of gene COL1A1 from GPT 3.5 using the prompt: *Please summarize the major function of a gene: COL1A1. Use academic language in one paragraph and include pathway information.*:

*COL1A1 is a gene that encodes the alpha1 chain of type I collagen,* ***which is the most abundant extracellular matrix protein and a major component of the bone, skin, tendon, and other connective tissues. The gene plays a crucial role in the synthesis and assembly of type I collagen fibers, contributing to the structural integrity and mechanical properties of these tissues. Additionally, type I collagen is involved in the regulation of signaling pathways, such as the TGF-$\beta$ signaling pathway, which plays a key role in the regulation of cell growth, differentiation, and extracellular matrix synthesis. Mutations in COL1A1 are associated with various connective tissue disorders, including osteogenesis imperfecta and Ehlers-Danlos syndrome. Therefore, understanding the function of COL1A1 is important in elucidating the pathogenesis of these disorders and developing potential therapeutic strategies.***

For the text description from NCBI, we need to clean the format of the original data and transfer them into text representations. Therefore, the sentences and words in this text are incoherent and their format does not strictly follow the grammar. Such difference poses the problem of model alignment with human value. Moreover, text descriptions of NCBI focus specifically on the variety of gene names, symbols, refseq status, and other properties. Moreover, it contained detailed functional information of the given gene. Some of the information may be redundant. One advantage of the text representation from NCBI is its authority, which means the reliability of text descriptions from NCBI is generally greater than the outputs from LLMs for researchers.

For the text description from GPT 3.5, we can see that the sentences and words are coherent and formalized under the correct grammar framework. This text focuses more on the overview of functions for this given gene, including its major functional tissues and cell types (for cell-type marker genes like CD79). Moreover, this text also includes the relation between COLA1A1 and certain diseases, highlighting the potential of this gene as a therapeutic target. Such prior information will be helpful in the in-silico treatment analysis research. Moreover, different prompts can generate different types of descriptions for the same gene, exploring the diversity of the outputs from LLMs is also an interesting research track.

Therefore, these two different types of text descriptions have their own advantages and disadvantages, which might be the explanation for their different suitable scenarios. One interesting research topic will be how to combine the advantages of these two kinds of text descriptions to enhance their downstream applications.

Here is an example for the text description of B cells from GPT 3.5 using the prompt: *Please summarize the major function of cell type: B. Use academic language in one paragraph.*:

*B cells, a crucial component of the adaptive immune system, play a pivotal role in* ***humoral immunity by generating antibodies (immunoglobulins) in response to the recognition of specific antigens. Following activation, B cells undergo clonal expansion and differentiation into plasma cells, which are specialized antibody-producing cells. The antibodies secreted by plasma cells serve as effectors in immune responses, neutralizing pathogens, facilitating their removal by phagocytic cells, and promoting various immune processes such as opsonization and complement activation. Additionally, B cells can differentiate into memory B cells, which confer long-term immunity by enabling a rapid and robust antibody response upon re-exposure to the same pathogen. Overall, B cells are central to the adaptive immune system's ability to recognize and respond to a wide array of infectious agents and contribute significantly to immune memory and protection.***

For the text description from GPT 3.5, we find that the text description is also coherent and follows the standard grammar. It summarizes the major functions of B cells as well as the cell-cell communication that B cells involve.

### B Analyzing mouse single-cell data.

In this section, we explored the contribution of LLM embeddings for analyzing omic data from Mouse. We first analyzed the ability of clustering in Figure 17 (a) for both single-cell data and spatial transcriptomic data. While gene embeddings from GPT 3.5 wa mode can generate clustering performance better than PCA for scRNA-seq data (MCA) [115], such embeddings cannot improve the clustering for spatial data sequenced by Slide-seq v2 [116]. Moreover, cell-type embeddings from GPT 3.5 can also carry information from Mouse, proved by high scores for evaluating clustering performance.

We also considered the cross-species analysis for the similarity of embeddings from Human genes and embeddings from Mouse genes, which can enrich the study of gene similarity in addition to orthologous genes. We computed the Euclidean distance (ED) for all gene pairs based on gene embeddings from Mouse and Human, shown in Figure 17 (b). We then ranked the ED to retrieve most similar genes. To verify the similarity, we figured out that the closet gene pairs were the orthologous genes, thus the gene embeddings from LLMs could also reflect the common information from two species.

### C Understanding the effect of the number of cells and the number of genes towards GenePT and scELMo.

In this section, we further investigated the difference between the two averaging modes and the application scenarios and analyzed the relationship between the attributes of raw data and clustering effects.

First, we directly plotted the visualization results of three different methods for the hPancreas-train dataset in Extended Data Figure 15. From this figure, we found that using neither gene embeddings from GenePT nor GPT 3.5 with the *aa* mode could preserve the cell-type-specific clusters in the space of UMAPs. However, using gene embeddings from GPT 3.5 with *wa* mode could preserve the major cell-type-specific clustering information. Based on this interesting observation, we further compared the weights used for these two averaging modes based on KL-divergence (KL-Div) [110], which could reflect the difference for the distribution of gene expression levels in each cell and a uniform distribution with $P = \frac{1}{m}$. For each dataset, the row sum is one for both these two cases. Based on Extended Data Figure 18 (a), we found that for all the four datasets we compared in the cell clustering task, the KL-div was larger than two and none of them had cells with zero divergence. Therefore, the distribution of gene expression levels carried more information compared with the weights based on uniform distribution. Moreover, we also showed that using *wa* mode was better for batch effect correction. Therefore, the *wa* mode is more suitable to handle tasks under the zero-shot learning framework.

Second, we analyzed the relation between the clustering performance and the number of genes. The first scenario we intended to investigate is the relation between the number of recorded genes and the clustering performance. In Extended Data Figure 18 (b), we display the change of clustering metrics with respect to the change of recorded genes based on the hPancreas-train dataset using *aa* mode. We noticed that the *wa* mode was not suitable for this research because we might have cells with zero expression by filtering some genes. There is no obvious correlation between the number of recorded genes and the clustering performance. Moreover, since the sources of GenePT or scELMo do not match all of the genes for every scRNA-seq dataset, sometimes we need to fill the gene embeddings of missing genes as zero. Therefore, we also investigated the relation between the number of matched genes and the clustering performance, shown in Extended Data Figure 18 (c). From this figure, we still did not observe a strong correlation between the number of matched genes and the clustering performance under the *aa* mode. However, for the *wa* mode, we found an obvious correlation between these two values. Therefore, having more matched genes can contribute to cell clustering under the *wa* mode of scELMo. Such conclusion also demonstrates the importance of extending our databases of feature embeddings.

Third, we analyzed the relation between the clustering performance and the number of cells. We subsampled different proportions of cells from the large-scale Onek1k PBMC dataset and computed the clustering results under different numbers of cells. Based on Extended Data Figure 18 (d), we found that there was no obvious correlation between the number of cells and the clustering performance for Onek1k PBMC dataset. Based on this dataset, we also explore the relationship between memory usage and cell numbers, shown in Extended Data Figure 19. The minimal requirement for loading the Onek1k PBMC dataset is 30 GB, and the growth is linear ($O(n)$ level). Therefore, cell number may not be a factor that can affect the performance of gene embeddings in this task. Moreover, scELMo is also capable of the analysis of large-scale scRNA-seq datasets.

### D Analysis of multi-omic data integration.

In this section, we explored the possibility of utilizing gene embeddings from GPT 3.5 to resolve multi-omic data integration task. Here we consider datasets from scRNA-seq and scATAC-seq without paired information. To reduce the dimensions of the scATAC-seq dataset, we transfer the feature information of such dataset from the space of peaks to the space of gene activity scores. The visualization results are summarized in Extended Data Figures 16 (a) and (b). According to these figures, we can still observe significant batch effect or the difference of cell embeddings from the cells with same cell types. Therefore, the function of scELMo for multi-omic data integration under the zero-shot learning framework is not good. Moreover, based on Extended Data Figure 16 (c), neither the *wa* mode nor the *aa* mode can improve the $S_{batch}$ score and the $s_{bio}$ score significantly. Incorporating the cell-type information into the cell embeddings space can significantly improve the averaged scores, but for metrics like iLISI to evaluate the mixture of batch information, such embeddings still had zero score. Therefore, scELMo is not capable of multi-omic data integration under the zero-shot learning framework.

### E The contribution of finetuned model in the in-silico treatment analysis.

In this section, we demonstrated the necessity of using a finetuned model rather than zero-shot learning for in-silico treatment analysis. In Extended Data Figure 20 (a), we display the change of CS for the same group of DEGs under the ascending aortic aneurysm disease and all genes were not significant for the Ascending only state. Moreover, the change of CS under PCA was nearly constant by varying different genes for removal, and the change of CS was very small for cell embeddings based on GenePT across all three states. Therefore, we concluded that the cell embeddings from either GenePT or PCA were not capable of modeling this disease without the knowledge of the intercellular variability due to diseases. Moreover, based on Extended Data Figures 20 (b) and (c), such variability was covered by the noise in the original expression space. Therefore, we need to learn a model that can distinguish the cells under different conditions as well as generate representative cell embeddings for the inference of novel therapeutic targets. Our ideas aligned with the strategy adopted by Geneformer.

Moreover, we compared the classification metrics for the cell-level disease condition under different gene embeddings for these two datasets and displayed the results in Extended Data Figure 21. From this figure, we found using gene embeddings containing information from genes generally had better performance than using embeddings from random numbers. Moreover, the classification results of scELMo based on gene embeddings from GPT 3.5 were slightly better than the results based on gene embeddings from GenePT. Therefore, using gene embeddings from GenePT and GPT 3.5 all contributed to generating representative latent space for different diseases.

We also considered genes whose silence might shift the cell embeddings from the control condition to diseased conditions. Therefore, our candidate genes became DEGs for the control case and we reversed our score to keep its direction (higher score means that the removal of this gene contributes to the change of cells from the healthy condition to diseased conditions). Our results are summarized in Extended Data Figures 22 (a) and (b). Based on our results, we identified different number of genes for different diseases. Moreover, there existed gene overlap across three states of the given disease, which implied that the removal or silence of such gene might have different contributions for different disease. Because the function of genes is closely related to the pathway [117], our findings can help analyze the pathogenesis of some diseases.

### F Supplementary figures

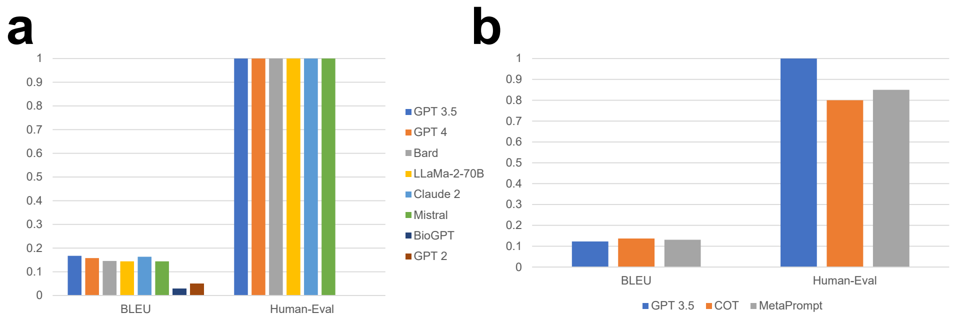


**Extended Data Fig. 1** Evaluations of descriptions of cell types and prompt engineering approaches. (a) Metrics for evaluating meaningful outputs of cell types across different LLMs. (b) Metrics for evaluating meaningful outputs of cell types across different prompt engineering approaches.

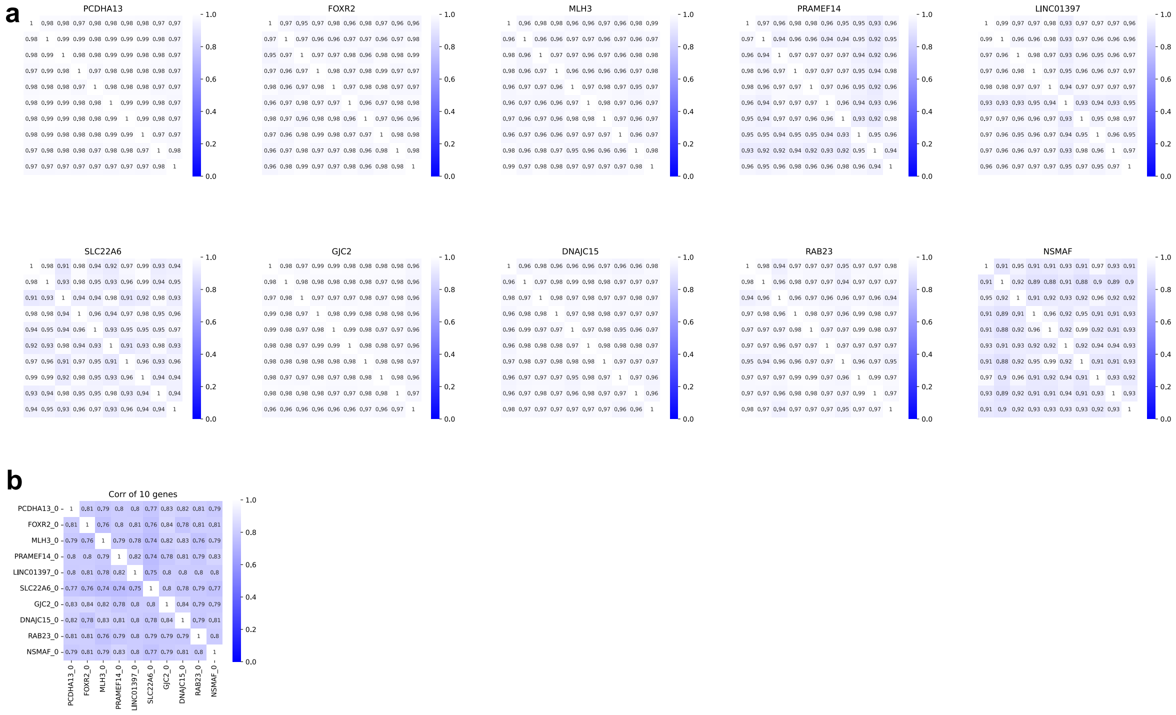


**Extended Data Fig. 2** Evaluations of the similarity for embeddings under different conditions. (a) Heatmaps of correlation of embeddings from the same gene under 10 different LLM outputs. We randomly selected 10 genes and generated the LLMs' outputs for these 10 genes based on the same set of prompts. We then computed the embeddings of these outputs and calculated the Pearson correlation for these embeddings, hence we have 10 different heatmaps to represent the results for 10 different genes. (b) Heatmap of the correlation of embeddings from 10 different genes. We computed the Pearson correlation for the embeddings of different genes. The number $_0$ represents the index of embeddings we computed based on 10 different LLM outputs.

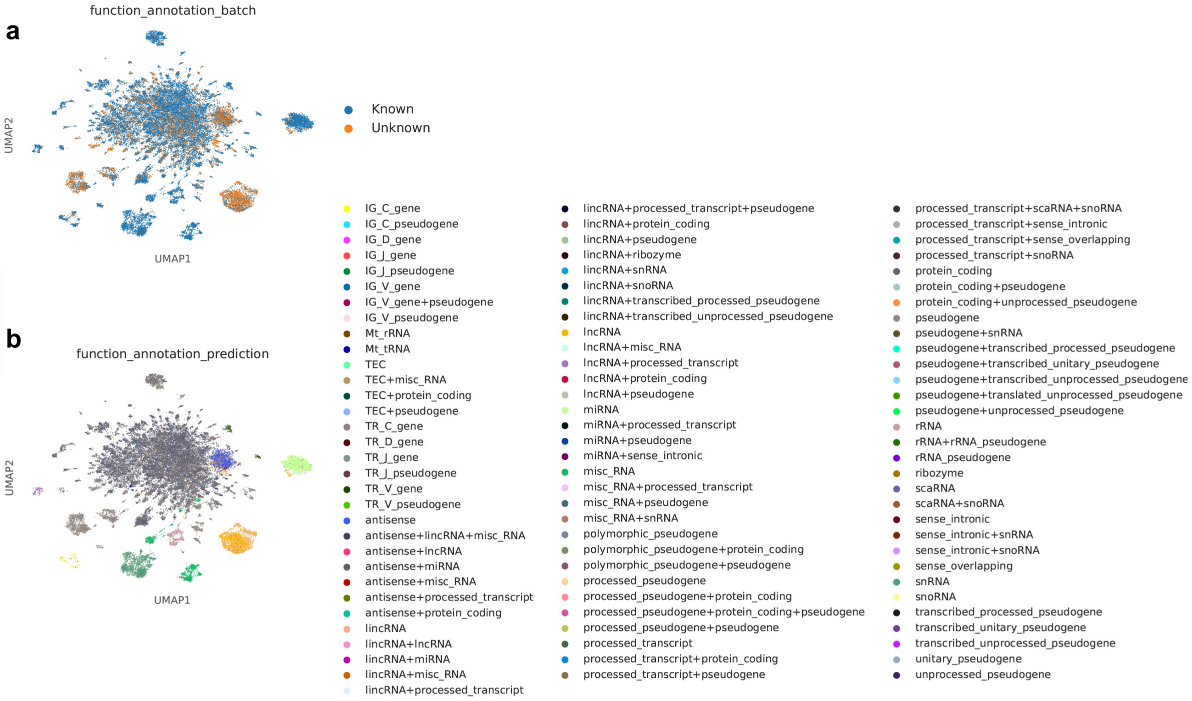


**Extended Data Fig. 3** UMAPs for the visualization of gene functional information. (a) UMAPs for the genes with known functional information and unknown functional information. (b) UMAPs for the genes with annotated functional information based on a kNN classifier. For genes with multiple functional annotation, we combined the functions as a new label.

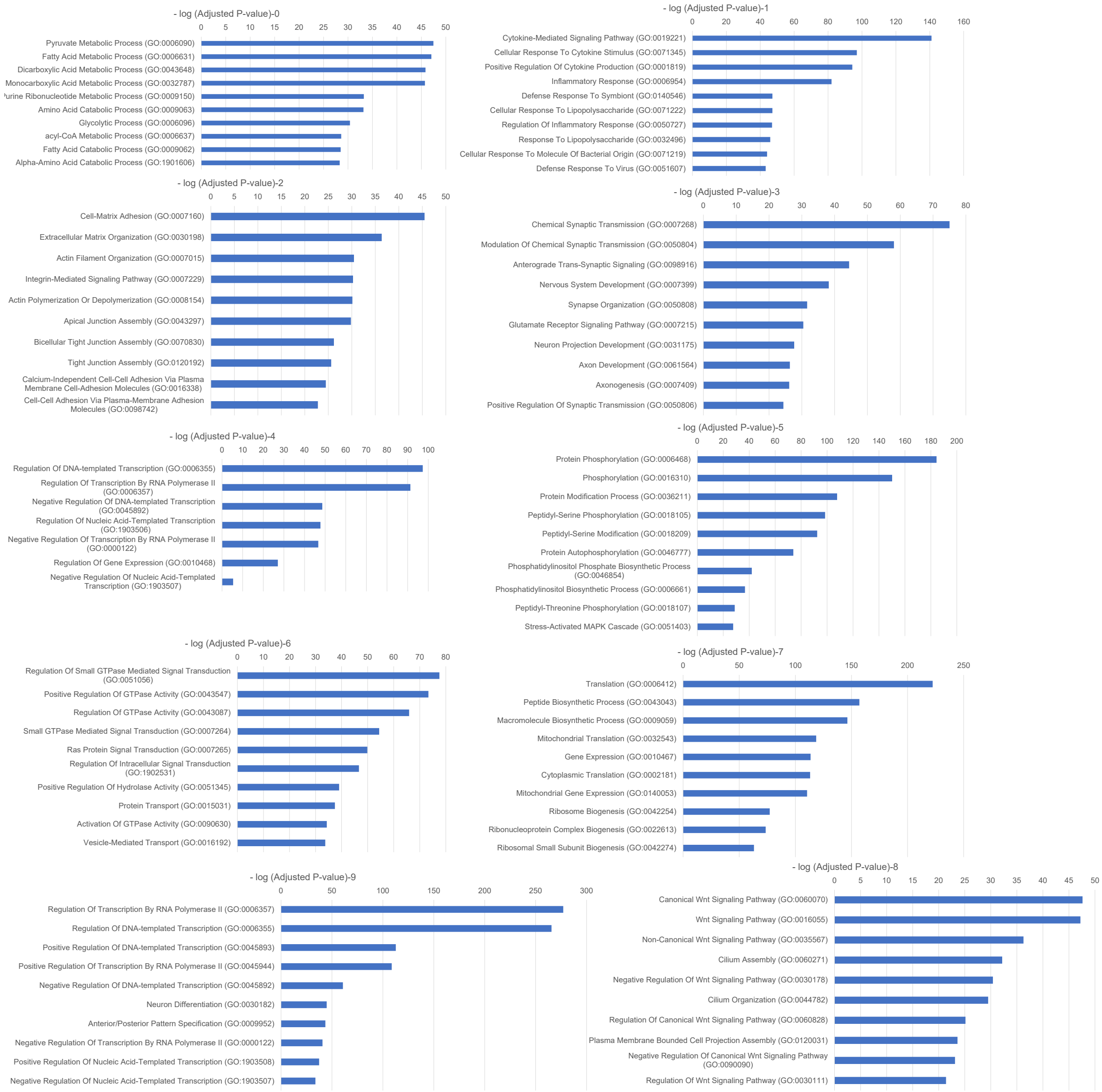


**Extended Data Fig. 4** Visualizations of gene pathway information for different clusters based on generated gene embeddings.

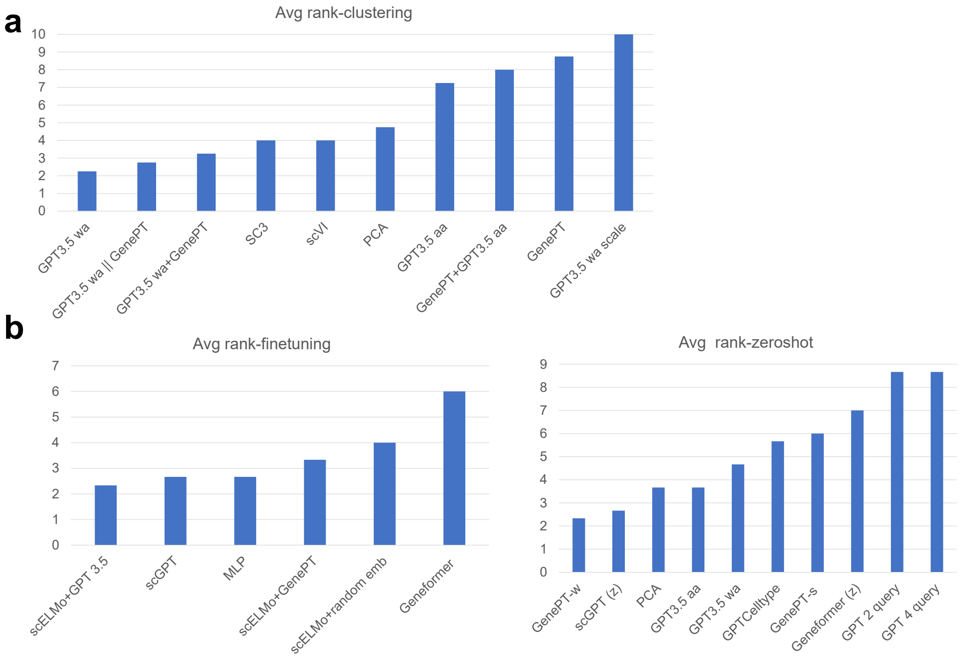


**Extended Data Fig. 5** Average ranks for clustering and cell-type annotation. (a) Average-rank information for different methods across datasets. (b) The left panel represents the average-rank information of methods based on fine-tuning for cell-type annotation. The right panel represents the average-rank information of methods based on zero-shot learning for cell-type annotation.

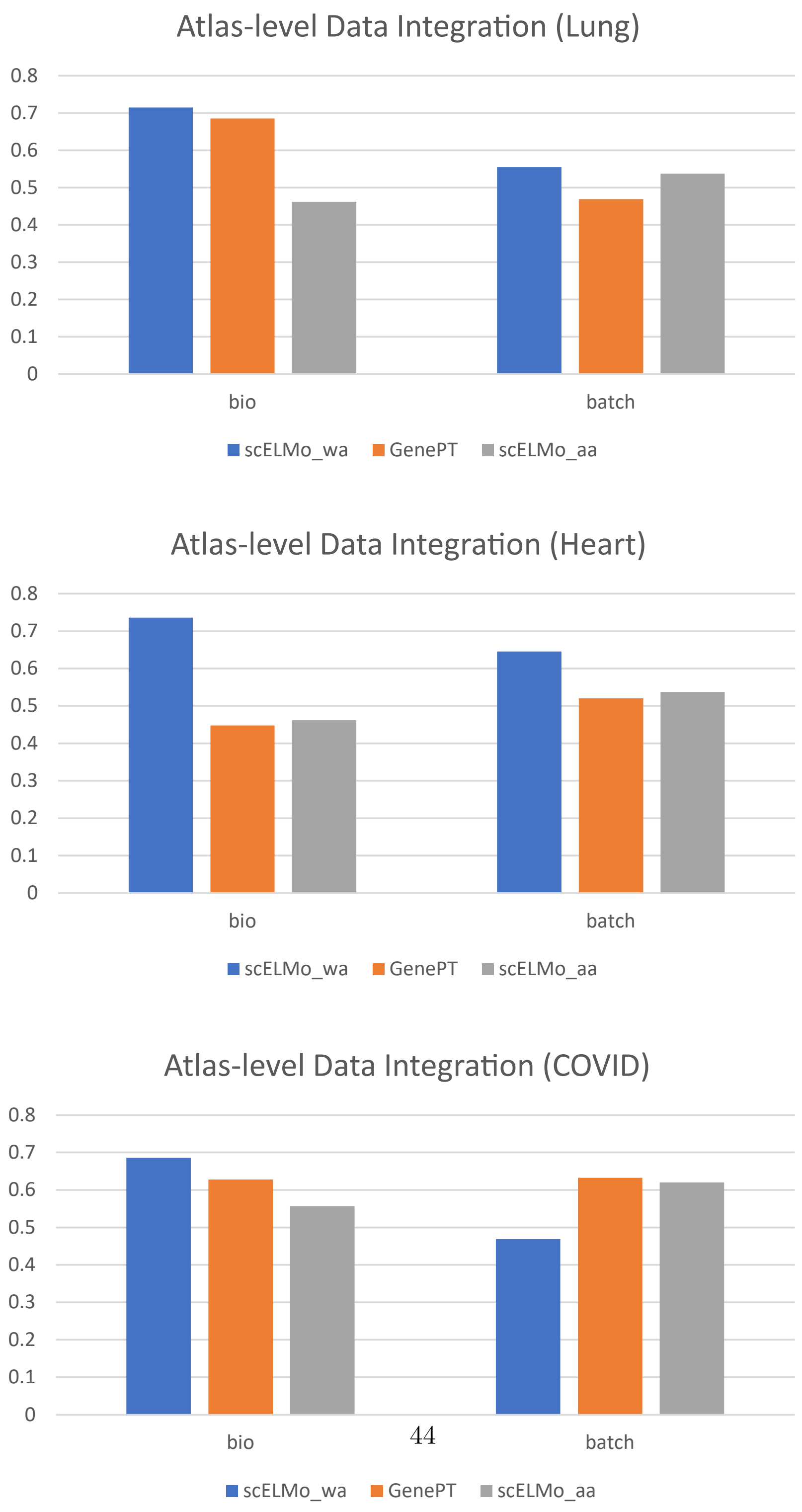




**Extended Data Fig. 6** Batch effect correction scores of scELMo and GenePT for atlas-level scRNA-seq dataset.

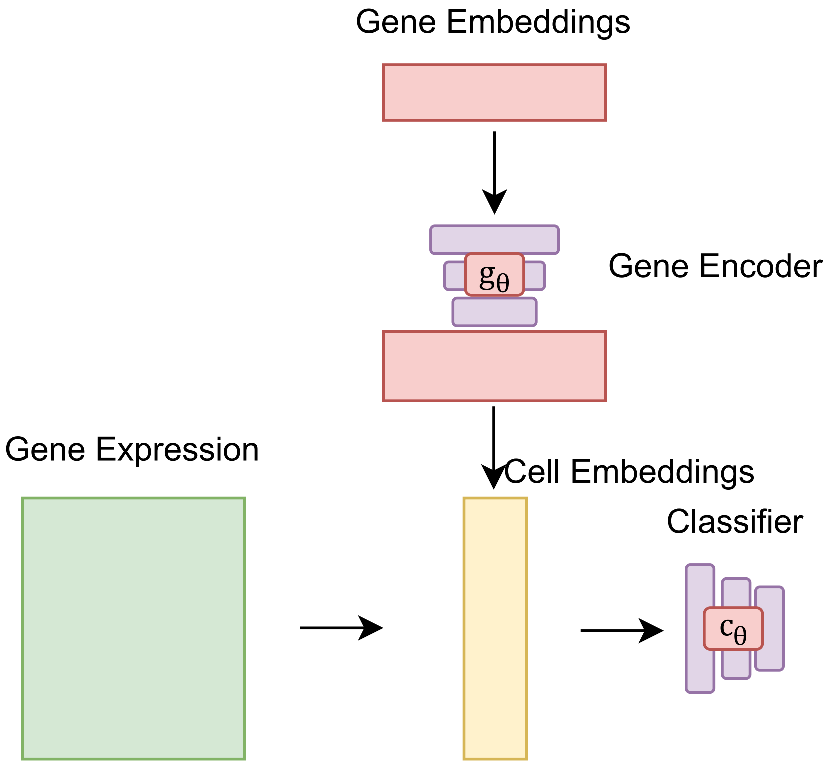


**Extended Data Fig. 7** Model architecture of scELMo for learning cell states or disease states.

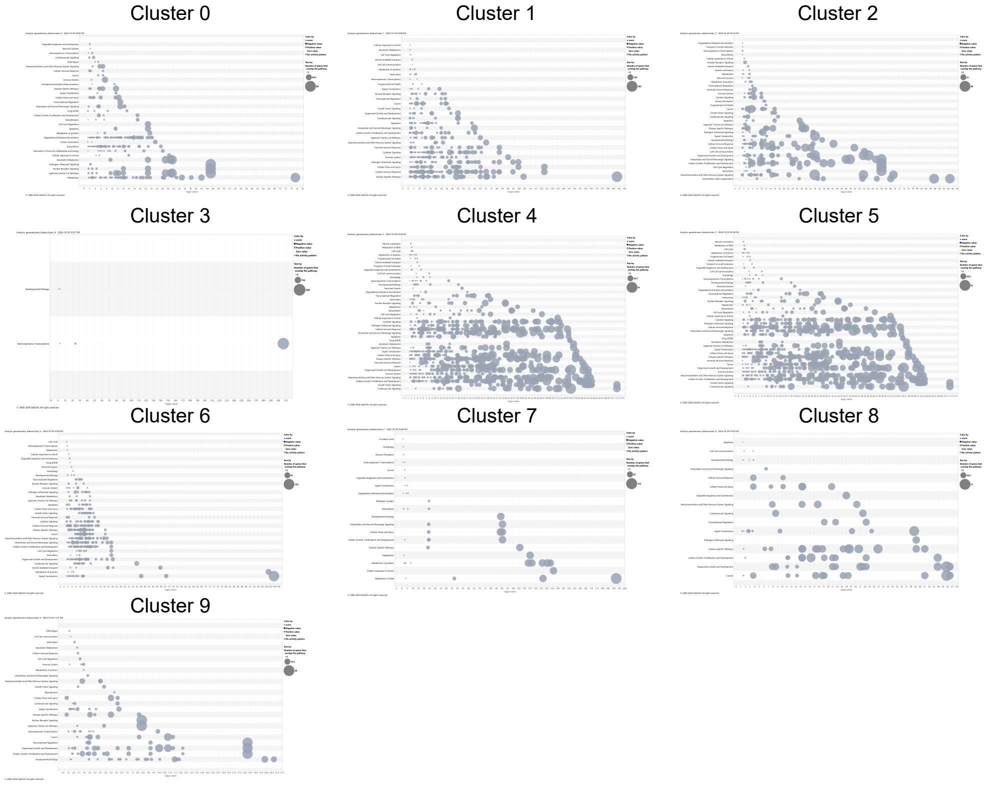


**Extended Data Fig. 8** Bubble plots for the pathway information discovered by IPA for each protein-encoding gene cluster. The size of each bubble represents the number of genes in the given pathway. The z-score value can be ignored as we do not incorporate gene expression information.

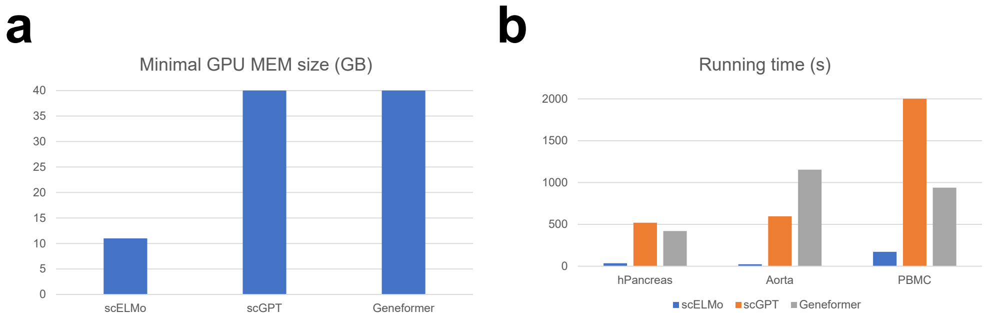


**Extended Data Fig. 9** Comparisons of resources. (a) The plot for minimal GPU memory requirements across different FMs. (b) The plot for running time of the cell-type annotation task across different FMs.

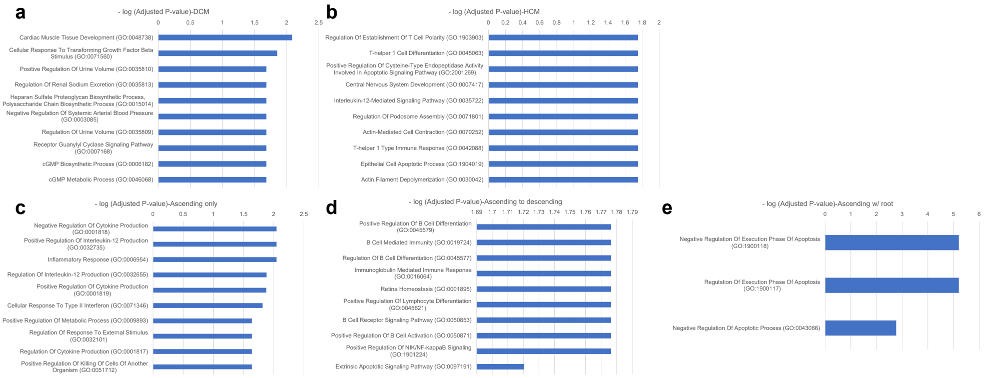


**Extended Data Fig. 10** Visualizations of gene pathway information for different conditions based on selected genes. (a) The GO enrichment results of target therapies for DCM. (b) The GO enrichment results of target therapies for HCM. (c) The GO enrichment results of target therapies for Ascending only. (d) The GO enrichment results of target therapies for Ascending to descending. (e) The GO enrichment results of target therapies for Ascending w/ root.

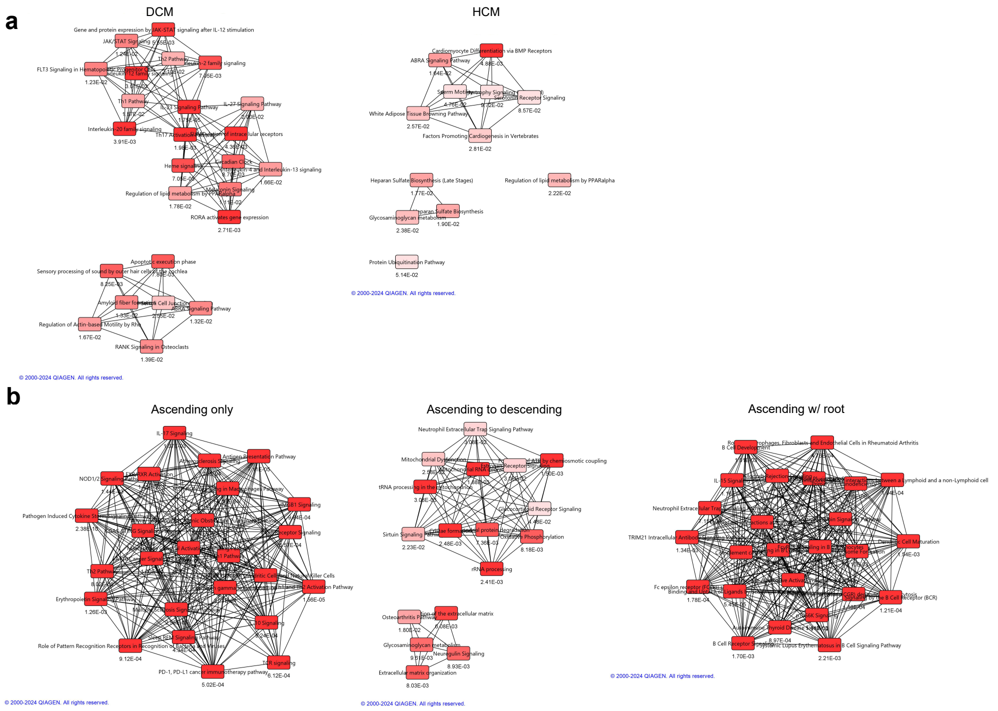


**Extended Data Fig. 11** Visualizations of IPA information for different conditions based on selected genes for in-silico treatment analysis. (a) The pathway information from the Heart dataset. (b) The pathway information from the Aorta dataset. The line represents pathway interaction, and we also show the p-value for each selected pathway.

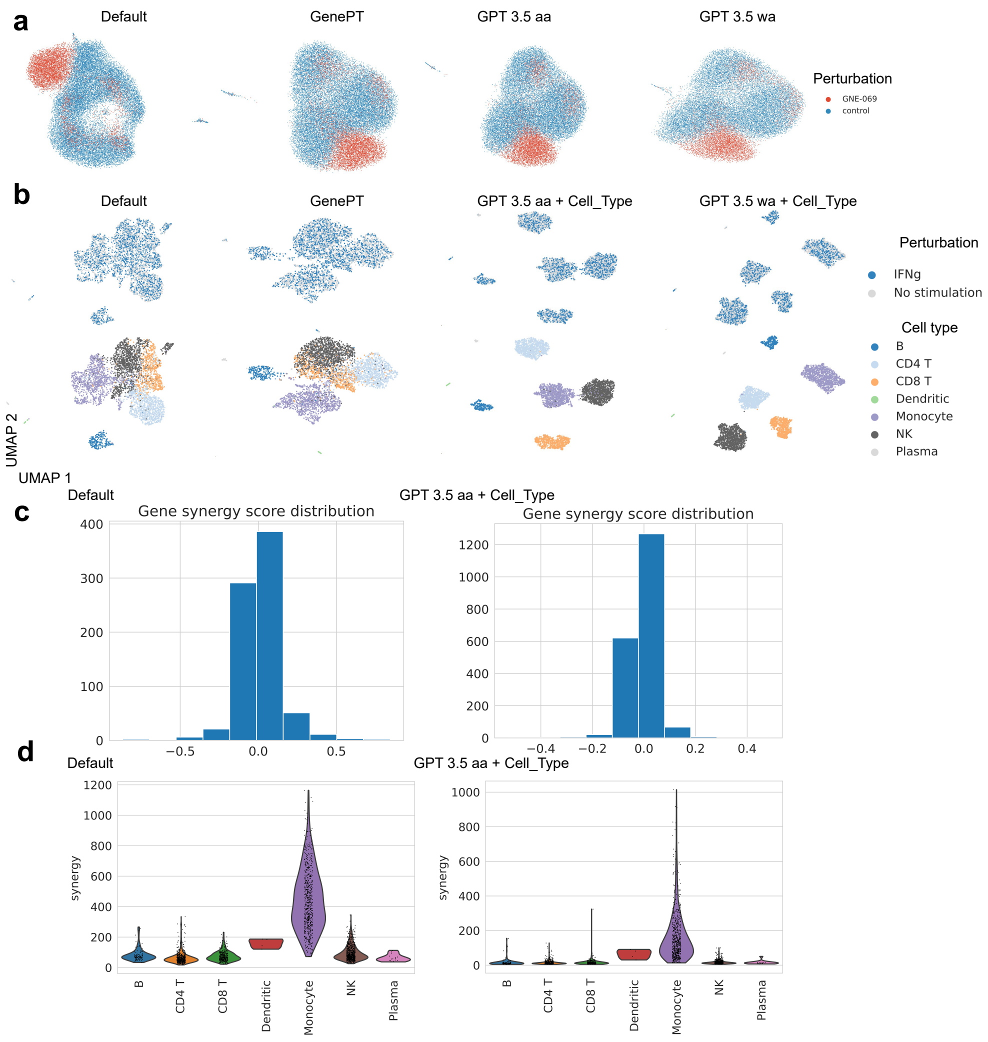


**Extended Data Fig. 12** UMAPs for the results of CINEMA-OT under different input settings and datasets. (a) UMAPs visualization for the confounder space of CINEMA-OT under different methods based on the ChangYe2021 dataset. All cells in this dataset have the same cell type. (b) UMAPs visualization for the confounder space of CINEMA-OT under different methods based on perturbed PBMC dataset. The labels for the UMAPs include perturbation conditions (upper panel) and cell types (lower panel). (c) Plots for the gene synergy score distribution, labelled by different methods. (d) Plots for the gene synergy distribution across cell types, labelled by different methods.

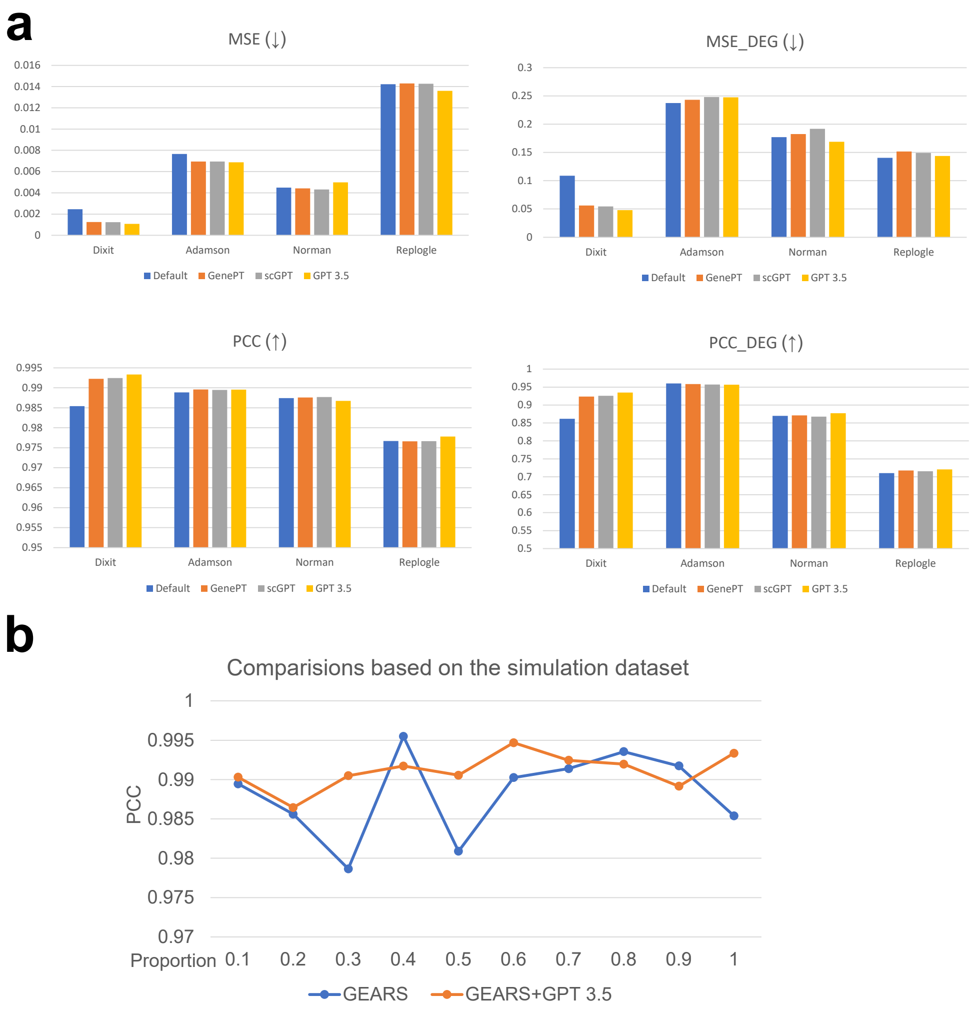


**Extended Data Fig. 13** The results of perturbation prediction for all datasets. (a) The MSE, MSE_DEG, PCC and PCC_DEG of all benchmarked methods across all datasets. The direction of the arrow represents the direction of better results. (b) Prediction results under the simulation dataset by subsetting the Dixit dataset with different cell-level proportions.

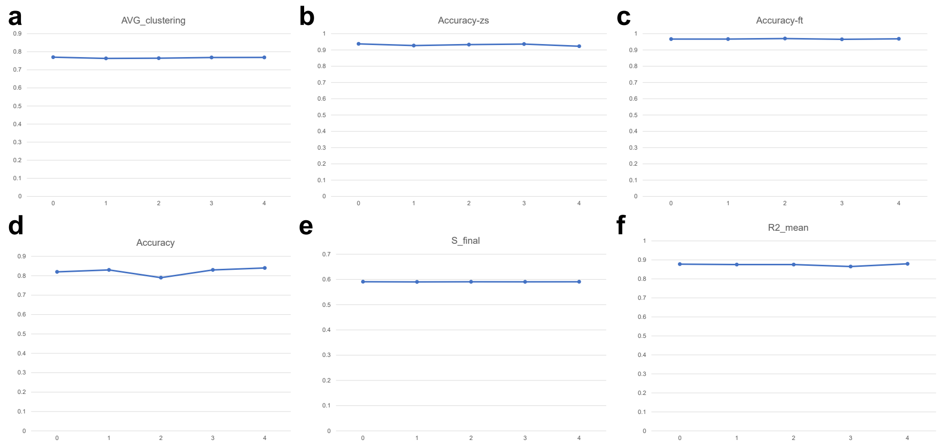


**Extended Data Fig. 14** Results of all downstream applications with gene embeddings from different random seeds 0-4. Here we used different random seeds to generate the descriptions of genes. (a) Results of clustering metric based on hPancreas-train dataset. (b) Results of classification metric based on the hPancreas dataset with zero-shot framework. (c) Results of classification metric based on the hPancreas dataset with fine-tuning framework. (d) Results of classification metric based on the Heart dataset for in-silico treatment analysis. (e) Results of integration metric based on the Cytof-CITE-seq dataset for batch effect correction. (f) Results of regression metric based on the CPA example dataset for perturbation prediction. We record the average R2 scores of each seed and display them.

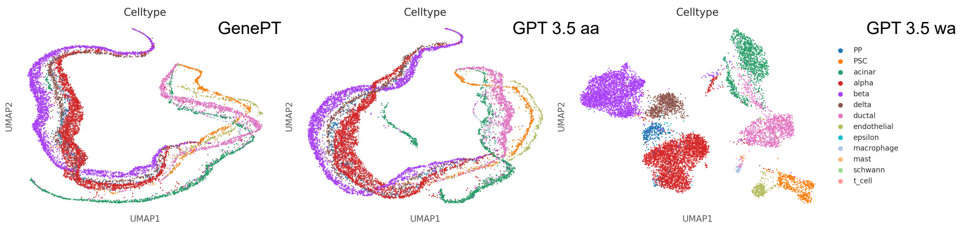


**Extended Data Fig. 15** UMAPs for cell embeddings with different sources based on the hPancreas-train dataset. Each panel is colored by the cell types. (a) UMAPs for the cell embeddings based on GenePT. (b) UMAPS for the cell embeddings based on *aa* mode. (c) UMAPs for the cell embeddings based on *wa* mode.

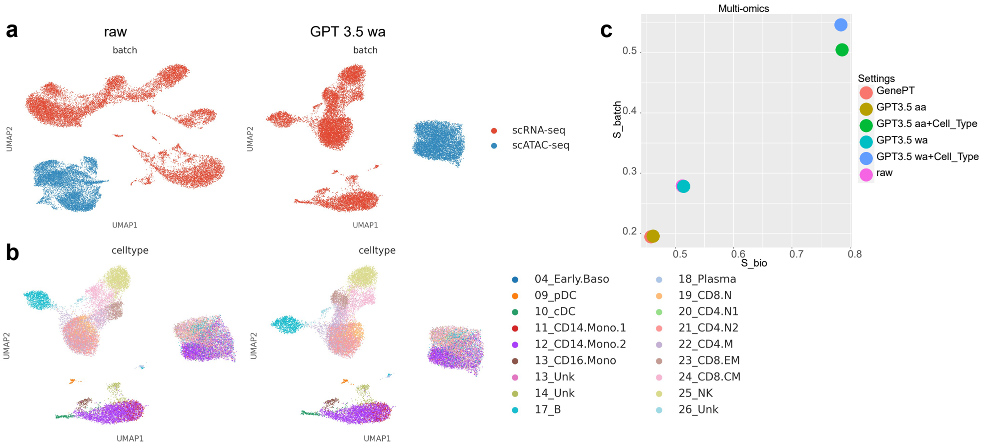


**Extended Data Fig. 16** Results of batch effect correction for multi-omic data (scATAC-seq, scRNA-seq). (a) UMAPs of batch information for the raw data and cell embeddings from GPT 3.5 *wa* mode. (b) UMAPs of cell-type information for the raw data and cell embeddings from GPT 3.5 *wa* mode. (c) Evaluations of the batch effect correction for multi-omic datasets across different methods.

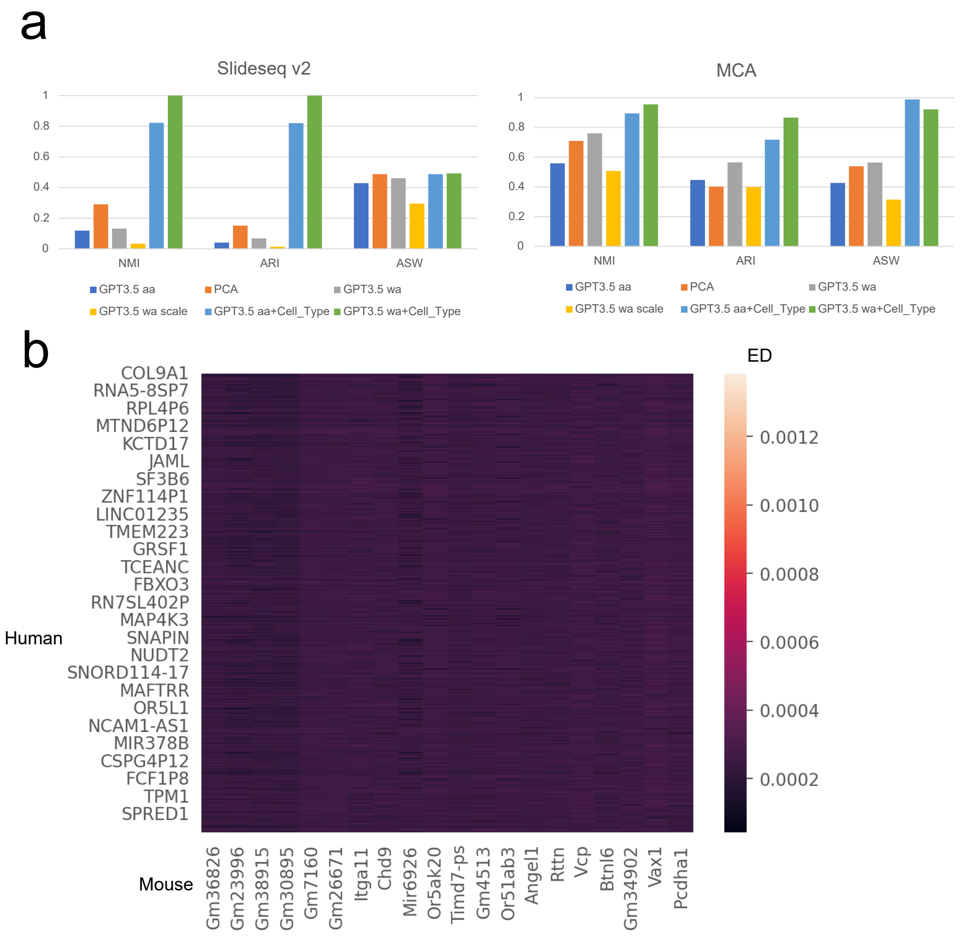


**Extended Data Fig. 17** Results of our exploration for gene embeddings from Mouse. (a) Clustering performance for Mouse data. The left panel represents the clustering metrics based on Slide-seq v2 data. The right panel represents the clustering metrics based on MCA data. (b) A heatmap for gene-gene interaction colored by the value of ED.

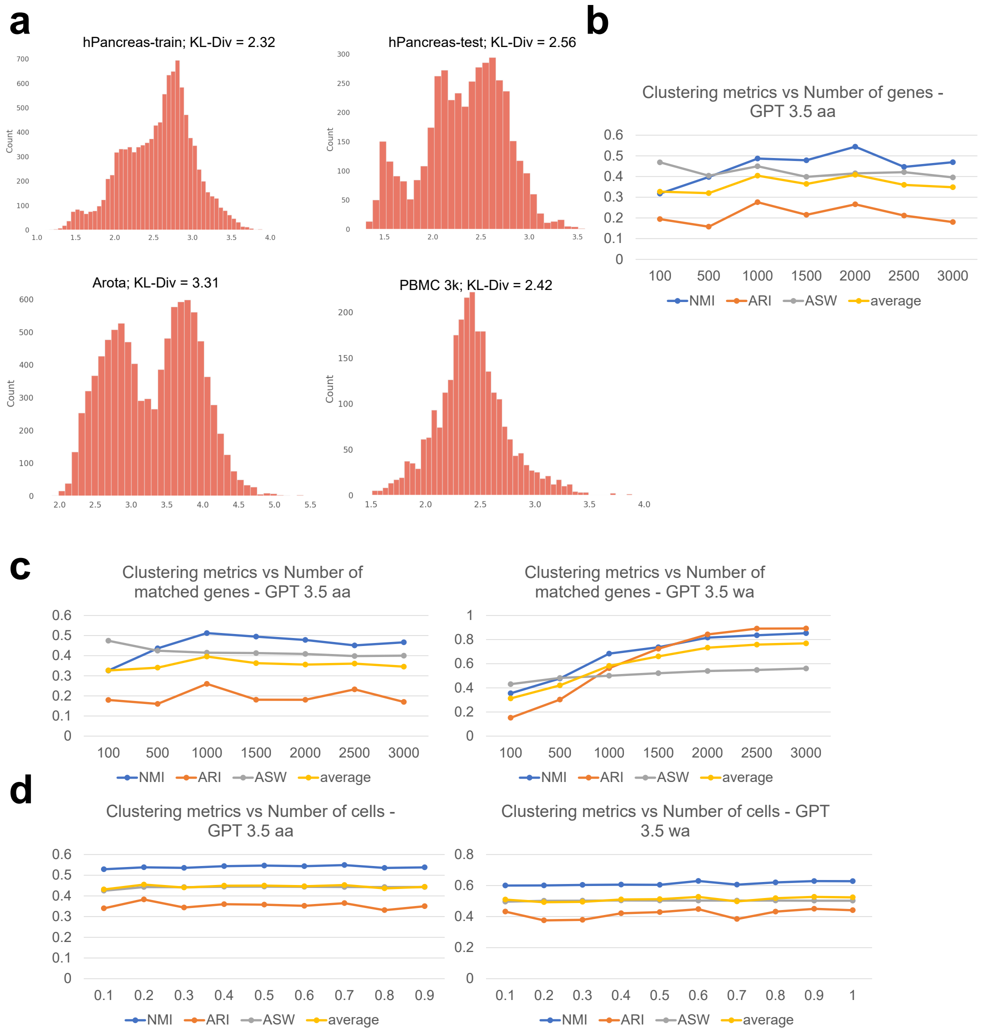


**Extended Data Fig. 18** Results of our exploration for the cell clustering task. (a) KL-Div for the two distributions across different datasets. (b) The relation between the number of recorded genes and clustering metrics is based on the GPT 3.5 *aa* mode. (c) The relation between the number of matched genes and clustering metrics is based on the GPT 3.5 *aa* mode (left panel) and the GPT 3.5 *wa* mode (right panel). (d) The relation between the proportion of cells and clustering metrics is based on the GPT 3.5 *aa* mode (left panel) and the GPT 3.5 *wa* mode (right panel).

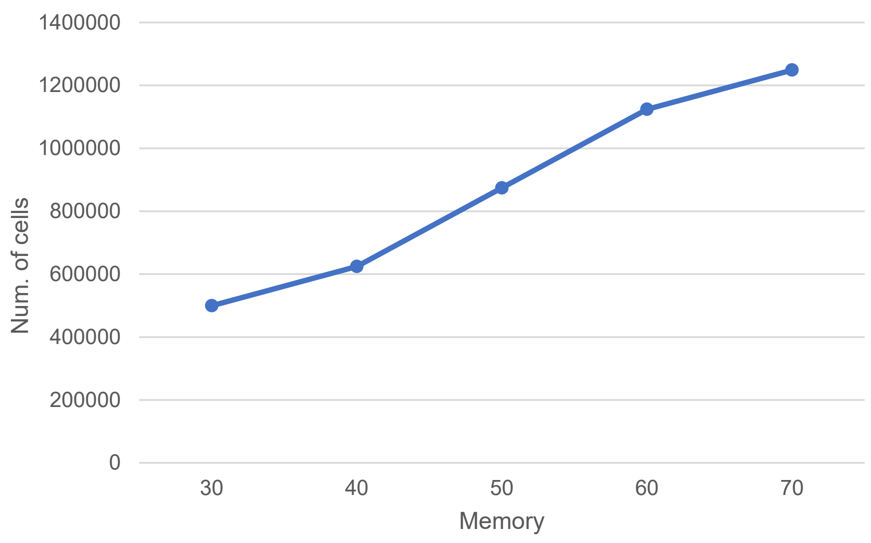


**Extended Data Fig. 19** The relationship between memory usage and its corresponding peak number of cells.

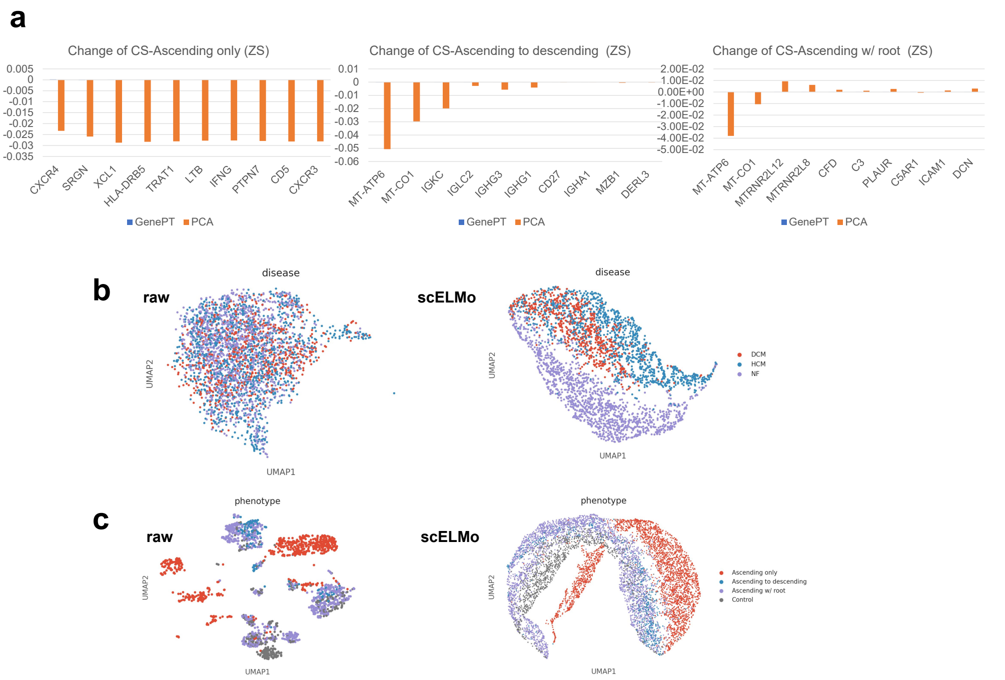


**Extended Data Fig. 20** Change of CS under the zero-shot (ZS) learning framework and UMAPs for visualization. (a) The change of CS based on cell embeddings from GenePT or PCA for the Aorta dataset. We considered all three different disease states. (b) UMAPs visualization for the original gene expression space (left panel) and cell embeddings from finetuned scELMo (right panel) based on the Heart dataset. Figures are colored by cell conditions. (c) UMAPs visualization for the original gene expression space (left panel) and cell embeddings from finetuned scELMo (right panel) based on the Aorta dataset. Figures are colored by cell conditions.

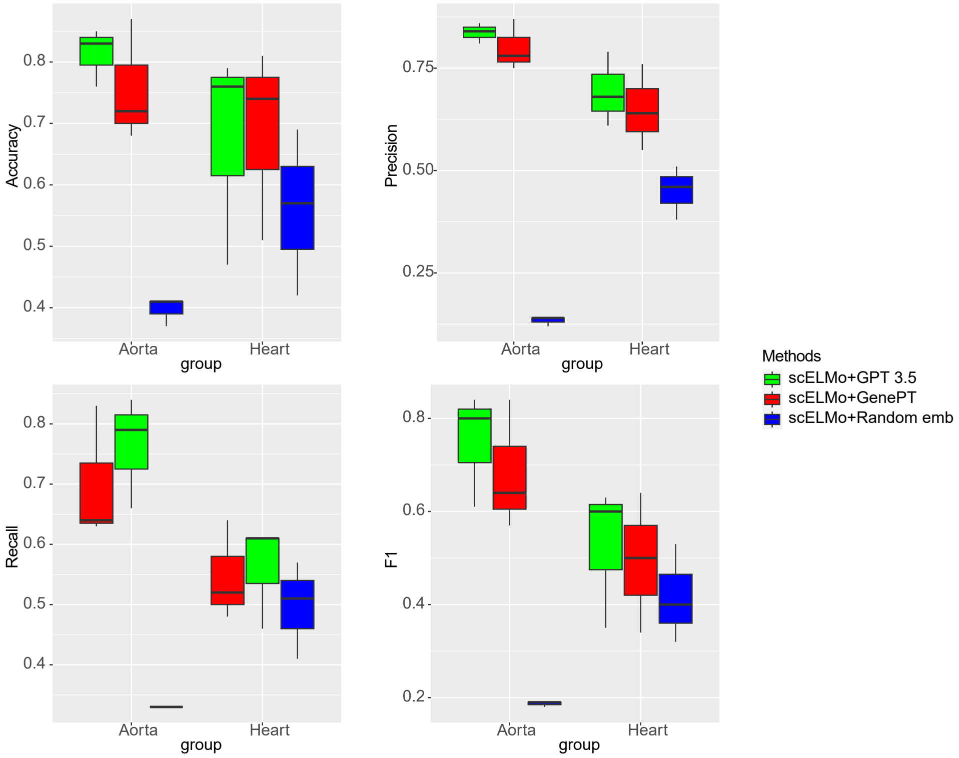


**Extended Data Fig. 21** Metrics for disease classification under different gene embeddings across the two datasets. Different panels represent values from different metrics, and we have four metrics in this task.

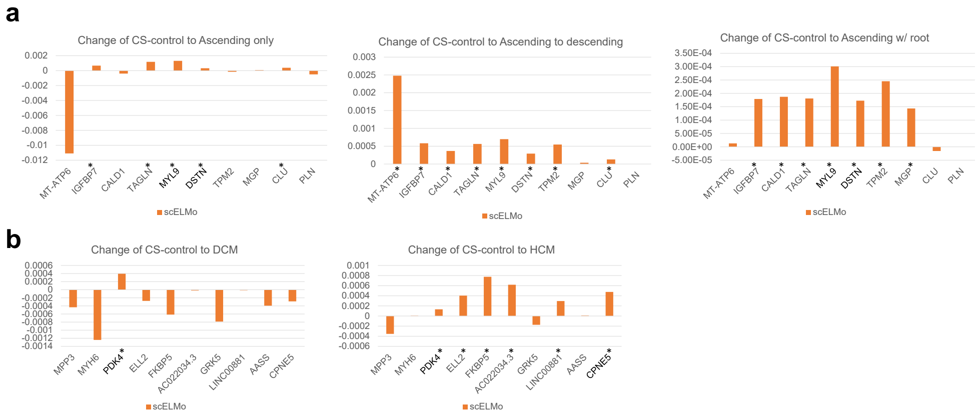


**Extended Data Fig. 22** Change of CS for silencing DEGs in the control case. (a) The change of CS based on cell embeddings from scELMo for the Aorta dataset. (b) The change of CS based on the cell embeddings from scELMo for the Heart dataset. We highlighted the genes detected by both GenePT and scELMo using stars (*) and marked the genes that were discovered by previous research as genes related to disease pathway using **bold** type.
